# Antibody ligation of carcinoembryonic antigen-related cell adhesion molecule 1 (CEACAM1), CEACAM3, and CEACAM6, differentially enhance the cytokine release of human neutrophils in responses to *Candida albicans*

**DOI:** 10.1101/2021.02.11.430790

**Authors:** Esther Klaile, Juan Pablo Prada Salcedo, Tilman E. Klassert, Matthias Besemer, Anne-Katrin Bothe, Adrian Durotin, Mario M. Müller, Verena Schmitt, Christian H. Luther, Marcus Dittrich, Bernhard B. Singer, Thomas Dandekar, Hortense Slevogt

## Abstract

Invasive candidiasis, often caused by *Candida albicans*, is an important healthcare-associated fungal infection that results in a high mortality rate of up to 40%. Neutrophils are the first line of defense during Candida infections. They can initiate various killing mechanisms and release cytokines to attract further immune cells to the site of infection. These responses are tightly controlled, since they can also lead to severe tissue/organ damage. We hypothesized that the regulation of *C. albicans*-specific neutrophil functions by the immunoregulatory *C. albicans* receptors CEACAM1, CEACAM3, and CEACAM6 are involved in the immune pathology of candidemia. Here, we analyzed the effects of the specific antibodies B3-17, 308/3-3, and 1H7-4B, respectively, targeting the three CEACAM receptors on *C. albicans*-induced neutrophil responses. We show that ligation of CEACAM6 significantly enhanced the response to *C. albicans*, as evidenced by the increased CXCL8/IL-8 secretion. By assessing the transcriptional responses, we found that CEACAM6 ligation and to some extent CEACAM1 ligation, but not CEACAM3 ligation, resulted in altered gene regulation of the *C. albicans*-stimulated neutrophils. Genes that were differentially regulated by the different CEACAM-targeting antibodies were analyzed for affected cellular processes and signaling pathways using various bioinformatics methods, including integrated network analyses and dynamic simulations of signaling cascades. Predicted changes in cellular pathways and cellular functions included CEACAM-specific alterations in apoptosis and cytokine secretion. In particular, we verified predicted changes in IL-1β/IL-6 expression in response to the antibody ligation of all three targeted CEACAMs and apoptosis induction by anti-CEACAM6 antibody treatment in presence of *C. albicans* stimulation. Specifically, CEACAM6 ligation by 1H7-4B enhanced neutrophil apoptosis and increased immediate and long-term cytokine release in responses to *C. albicans*. CEACAM1 ligation by B3-17 mainly enhanced the immediate secretion of pro-inflammatory cytokines, and CEACAM3 ligation by 308/3-3 increased the long-term release of pro-inflammatory cytokines. Taken together, we demonstrated for the first time that CEACAM receptors have an important and differential impact on the regulation of *C. albicans*-induced immune functions in human neutrophils.

## Introduction

Invasive candidiasis is the most common fungal disease among hospitalized patients in the developed world, and the fourth most common bloodstream infection in intensive care units (1). Invasive candidiasis is a major threat to immunosuppressed or critically ill patients and results in mortality rates as high as 40%, even when patients receive antifungal therapy; *Candida albicans* causes the majority of these infections (1).

Neutrophils are key players in the host defense against fungal pathogens. These short-lived innate immune cells react instantly and launch intracellular and extracellular killing mechanisms like the production of reactive oxygen species and microbicidal peptides, or the release of neutrophil extracellular traps (2). Neutrophils also release stored cytokines and produce new ones to attract further immune cells to the site of infection (3). While the total amounts of the secreted cytokines per neutrophil are rather small compared to other immune cells, neutrophils usually greatly outnumber mononuclear leukocytes in inflammatory sites by 1-2 orders of magnitude, implying that they can certainly have a major influence on the local and systemic long-term development of the immune response (3).

The neutrophil responses necessary for the quick eradication of pathogens are tightly controlled, since they can also have cytotoxic side effects and result in severe tissue/organ damage, that can subsequently lead to organ failure(s) and the death of the patient (2, 4, 5). Mechanisms for restricted neutrophil activity include the up or down regulation of various surface receptors and intracellular proteins within important signaling pathways, as recently comprehensively reviewed (6). Carcinoembryonic antigen-related cell adhesion molecules (CEACAMs) are immuno-regulatory surface receptors that are able to modulate the activity of a number of important immune receptors, including pattern recognition receptors Toll-like receptor 2 (TLR2) and TLR4 (7–11).

Three members of the CEACAM receptor family that recognize *C. albicans* surface structures are co-expressed on human neutrophils: CEACAM1, CEACAM3, and CEACAM6 (12, 13). The neutrophil marker CEACAM8 (CD66b) does not bind to *C. albicans* (12). While the extracellular domains of CEACAM receptors are composed of highly conserved Ig-like domains, their signaling properties differ widely (11, 14) and can be activating or inhibitory. CEACAM1 is also expressed on various other leukocytes and epithelial cell types (11, 15). The cytoplasmic domain of CEACAM1 bears an immunoreceptor tyrosine-based inhibition motif (ITIM) and recruits Src family kinases and SHP1 and SHP2 phosphatases. Importantly, it can act as both, an activating or an inhibiting receptor (8, 10, 12, 16–19) and delays apoptosis in granulocytes and lymphocytes (20–22). CEACAM3 is exclusively found on granulocytes and acts as an activating receptor via its immunoreceptor tyrosine-based activation motif (ITAM) upon ligation by bacterial pathogens (23, 24). CEACAM6 is GPI-anchored and constitutively localized to membrane microdomains, as demonstrated in transfected 293T cells (14). The modulatory roles of CEACAM1, CEACAM3, and CEACAM6 in cellular responses to bacterial pathogens were recently analyzed in detail (8, 25–31). In particular, a comprehensive study on the CEACAM-specific neutrophil responses to *Neisseria gonorrhoeae* in humanized murine neutrophil cell models shows that CEACAM3 mainly mediates neutrophil activation and bacterial killing, while CEACAM1 and CEACAM6 mostly mediate pathogen binding and phagocytosis (28). Interestingly, this study also demonstrates that the co-expression of CEACAM1 and CEACAM6 potentiate, rather than hinder, CEACAM3-dependent responses of neutrophils to the bacterial pathogen (28). Similar results were obtained for *Helicobacter pylori* in murine neutrophils from humanized transgenic mice, where human CEACAM1 expression was sufficient for the binding and the phagocytosis of this pathogen, but an increased ROS production and an enhanced CCL3 secretion was only found in human CEACAM3/CEACAM6-expressing neutrophils, independent of the additional presence of human CEACAM1 (31). However, other studies using neutrophils from Ceacam1^-/-^ mice describe CEACAM1 as an important inhibitory receptor that negatively controls granulopoiesis as well as the LPS-induced IL-1β and ROS production in neutrophils (10, 32). The same mice were used in a study demonstrating the CEACAM1-mediated negative regulation of MMP9 release by mouse neutrophils (12).

In the present study, we used CEACAM mono-specific monoclonal antibodies to ligate CEACAM1, CEACAM3 and CEACAM6, respectively, on human neutrophils to analyze their effects on *C. albicans*-induced responses. Recent studies using different monoclonal antibodies against CEACAM receptors in the absence of pathogens demonstrate that the ligation of either CEACAM on neutrophils resulted in signaling events, priming, and in an enhanced β2-integrin-dependent adhesion to endothelial cells (33, 34). We assessed in the present study, whether neutrophil treatment with various CEACAM-specific antibodies alters different aspects of the immediate response (30-120 min) of human neutrophils to *C. albicans*, including CXCL8 secretion, ROS production, and transcriptional responses. Moreover, systems biological analysis of the neutrophil transcriptional responses to the CEACAM-specific antibodies in presence or absence of *C. albicans* revealed affected cellular functions that were verified for their long-term (4-24 h) biological effects *in vitro*, like apoptosis and the *de novo* production of IL-6 and IL-1β.

## Results

### Increased *C. albicans*-induced CXCL8 release by neutrophil pretreatment with anti-CEACAM6 antibody

In order to establish an easy read-out for the activation of human neutrophils by *C. albicans*, we first ascertained that this stimulation resulted in the early secretion of CXCL8 (also known as IL-8; Fig 1A). Indeed, stimulation with this fungal pathogen resulted in the rapid secretion of approximately 600 pg CXCL8/ml within 2h (10^7^ neutrophils/ml). Pretreatment with the mouse IgG1 isotype control antibody MOPC-21 did not alter CXCL8 release in response to *C. albicans* (Fig 1A). We then used different mouse monoclonal IgG1 antibodies either monospecific for CEACAM1, CEACAM3, and CEACAM6, or cross-reactive to more than one CEACAM receptor, (as indicated in Fig 1B), respectively, for neutrophil ligation before *C. albicans* stimulation. Treatment with the CEACAM6-specific monoclonal antibody 1H7-4B resulted in a strong and significant increase in CXCL8 release upon stimulation with *C. albicans*. One CEACAM1-specific antibody (B3-17) also displayed a considerable increase of CXCL8 release in presence of *C. albicans*, but results from B3-17 treatments did not reach statistical significance due to the high donor-specific variance for the CXCL8 responses. CEACAM6 ligation with 13H10 antibody increased CXCL8 secretion slightly but not statistically significant, while a recombinant CEACAM6-Fc protein that can also bind to CEACAM1 and CEACAM6 did not show any effect on CXCL8 secretion. CEACAM3 ligation with the specific 308/3-3 antibody did not significantly alter the CXCL8 response to *C. albicans* stimulation, while a slight but not significant increase was detected when monoclonal antibody 18/20 was used that cross-reacts with CEACAM1 and CEACAM3. None of the antibodies tested resulted in a reduction of *C. albicans*-induced CXCL8 levels.

**Fig 1:**
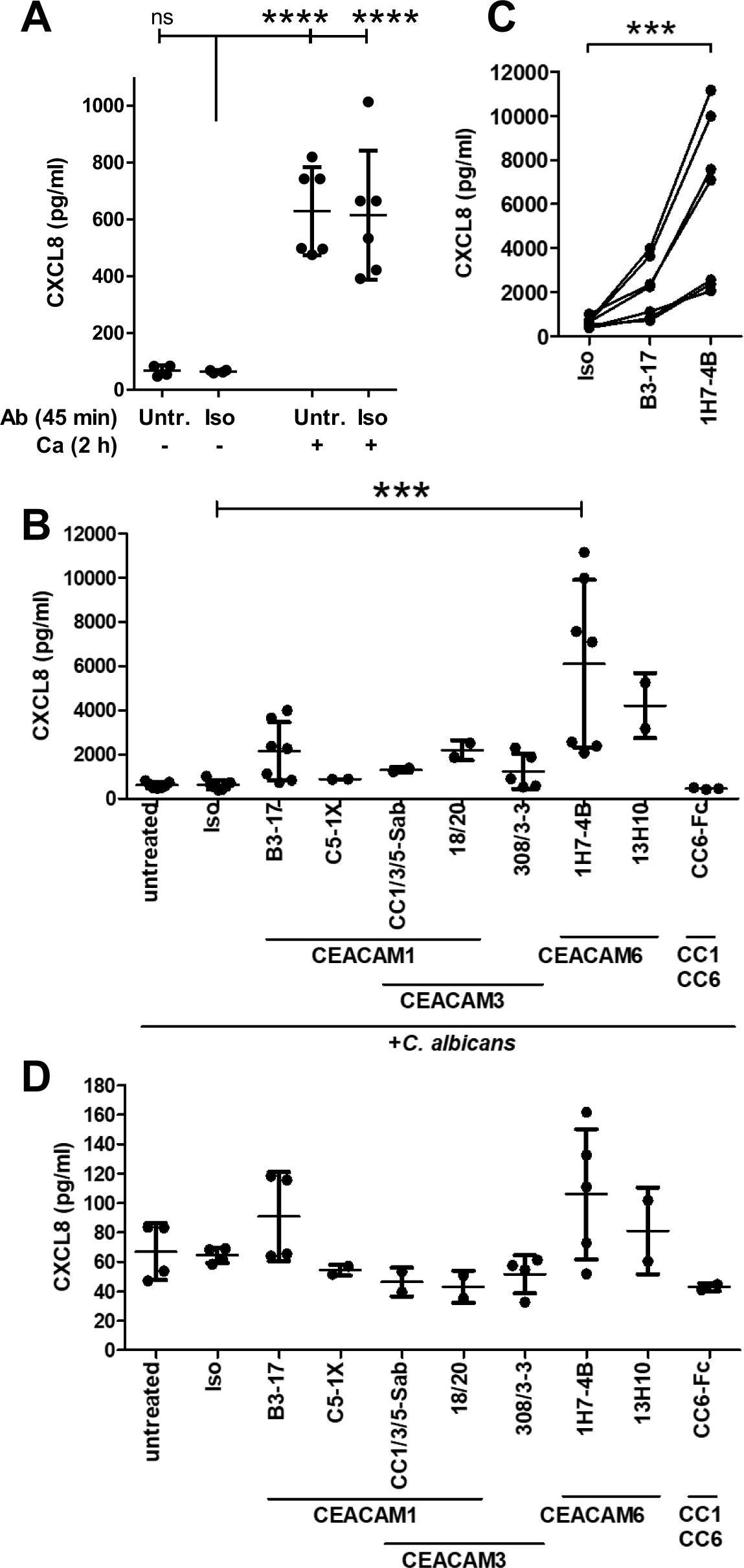
Altered *C. albicans*-induced CXCL8 release by neutrophil treatment with anti-CEACAM6 antibody. Human neutrophils (10^7^/ml) were left untreated or were incubated with 10 µg/ml mouse IgG1 isotype control antibody (clone MOPC-21), or monoclonal CEACAM-specific mouse monoclonal IgG1 antibodies for 45 min and were consecutively incubated with or without live *C. albicans* yeast cells (MOI 1) for 2 h. Antibody clones and their specific (cross-) reactivities for CEACAM1, CEACAM3 and CEACAM6 are indicated in (B) and (D). Cell culture supernatants were harvested and analyzed for CXCL8 concentrations in 2-7 independent experiments for the different treatment groups, represented by the single dots in the graphs (note that not all antibodies were tested in parallel). (A) CXCL8 release in untreated and isotype-treated neutrophils without and with *C. albicans* stimulation. (B) CXCL8 release in *C. albicans*-stimulated cells including all anti-CEACAM antibody treatments. Note t(C) Samples from six donors, also shown in (B), which were pre-treated with isotype control antibody, B3-17 and 1H7-4B, respectively, were plotted with linked samples from the same donor. Note the donor-independent relative increase of the CXCL8 response to *C. albicans* treatment after pre-stimulation with the CEACAM1- and the CEACAM6-monospecific antibody, respectively. (D) CXCL8 release after antibody treatments in absence of *C. albicans*. Statistical analyses were performed by One-Way ANOVA with Bonferroni post-test. ***p<0.005, ****p<0.001

Fig 1C highlights the CXCL8 responses in neutrophil cell culture supernatants from the same donors treated with IgG isotype antibody, B3-17, or 1H7-4B, respectively (same samples as displayed in Fig 1B). Here, differentially treated samples from the same donor are connected by lines. As depicted, B3-17 treatment always resulted in a small increase of the CXCL8 release compared to the isotype treatment, and 1H7-4B treatment resulted in a strong increase of CXCL8 release (up to 10-fold), illustrating the donor-independent relative effect of the different antibodies. A high variation in the release of CXCL8 and other cytokines by neutrophils from different donors are also found in other studies using different stimuli, including the CD11b/CD18 ligand Thy-1, LPS, and fMLP, and many studies therefore give the standard error instead of the standard deviation (35–38). Since the donor dependency of the strength of the induced cytokine release is inherent to this primary cell type, it cannot be avoided.

In absence of *C. albicans* stimulation, no significant differences in CXCL8 secretion after any antibody treatment were found (Fig 1D). However, in single neutrophil preparations, the three most effective antibodies in the presence of *C. albicans* (B3-17, 1H7-4B and 13H10) also resulted in a minor increase (max. 2-fold) of basic CXCL8 secretion when used alone in the absence of *C. albicans* (Fig 1D). The differences in CXCL8 concentrations in the supernatants between samples with antibody treatment in absence of *C. albicans* stimulation (30-170 pg/ml), and with antibody treatment in presence of *C. albicans* stimulation (400-11,000 pg/ml) ranged between one and two orders of magnitude, depending on the donor. For the following experiments, the CEACAM1-specific antibody B3-17 (“CC1”), the CEACAM3-specific antibody 308/3-3 (“CC3”), and the CEACAM6-specific antibody 1H7-4B (“CC6”) were chosen for further investigation.

### No influence of anti-CEACAM antibodies on neutrophil-mediated killing of *C. albicans*

We next analyzed the influence of the three CEACAM1/3/6-targeting antibodies on different cellular functions elicited directly upon the engagement of the neutrophil by the fungal pathogen. First, we tested if the antibodies affected *C. albicans* binding to neutrophil cell surfaces (S1 Fig). In accordance with our findings that the CEACAM family receptors are not critical for *C. albicans* adhesion to different epithelial cell lines (12) and CEACAM1-transgenic mouse neutrophils (39), neutrophil binding to fungal cells was not affected by antibody-mediated CEACAM ligation in two different donors (S1 Fig). A major killing mechanism employed by neutrophils is the oxidative burst. We therefore tested the production of reactive oxygen species (ROS) under the different antibody treatments in the presence or absence of *C. albicans* stimulation in two different donors (S2 Fig). While ROS production by *C. albicans* stimulation was clearly detected, none of the three antibodies used for ligation changed neither the basic ROS levels in absence of *C. albicans* nor the *C. albicans*-induced ROS levels after infection (S2 Fig). To test for effects of the antibody treatments on the overall efficiency of fungal killing by the neutrophils, we next analyzed the number of surviving fungal cells after 30 min co-incubation with the neutrophils. None of the tested CEACAM1-, CEACAM3- or CEACAM6-specific monoclonal antibodies showed a detectable effect on neutrophil killing efficiency (S3 Fig; about 10% of the fungal cells remained viable after 30 min incubation with human neutrophils at a 1:1 ratio, allowing for the detection of increased and decreased killing efficiencies, respectively; N=3).

### The *C. albicans*-induced differential transcriptional responses in neutrophils are affected by anti-CEACAM1 and anti-CEACAM6 ligation

CEACAM receptor ligation by various antibodies results in neutrophil priming and enhanced binding to endothelial cells (33, 34). Therefore, we next examined if antibody-mediated CEACAM ligation on neutrophil surfaces with B3-17 (“CC1”), 308/3-3 (“CC3”), and 1H7-4B (“CC6”) would also affect neutrophil transcriptional responses in the presence and the absence of *C. albicans* stimulation, respectively. Principal component analyses (PCA, S4 Fig) demonstrated that the different antibody treatments in the presence or the absence of *C. albicans* stimulation were well separated and that the data were of a good quality. The PCA also revealed that CC6 treatment had the largest influence on the neutrophil transcriptomic response induced by *C. albicans*, followed by CC1 treatment (S4 Fig). CC3 treatment before *C. albicans* infection only resulted in minor alterations in gene transcription. The three antibodies displayed similar hierarchies in absence of *C. albicans* stimulation (S4 Fig). The IgG treatment did not alter the transcriptional activity neither in the presence nor in the absence of *C. albicans* stimulation (S4 Fig), and we refer to these controls for all comparisons with anti-CEACAM antibody treatments.

We first analyzed the transcriptional response induced by *C. albicans* stimulation (S2 Table). IgG treatment with subsequent *C. albicans* stimulation of neutrophils resulted in 442 significantly differentially expressed genes (DEGs) compared to IgG treatment in absence of *C. albicans* stimulation: 213 DEGs were up-regulated and 230 were down-regulated (fold-change >±2, adjusted p-value <0.05; see S2 Table for details). GO enrichment analysis showed that the altered transcripts mostly comprised genes relevant to transcription or the regulation of various features important for the immune response, including cell migration and adhesion, the production of cytokines, neutrophil apoptosis and different corresponding signaling pathways, including MAP kinase signaling and NFκB signaling (S9 Table). These *C. albicans*-induced alterations in the transcriptional profile of human neutrophils after 2 h of stimulation were similar to those detected in other studies after 1 h of stimulation (40). Since this study aims to shed light on the role of CEACAM receptors in the neutrophil response to *C. albicans*, we investigated significant differences in the gene expression induced by the pre-treatment with the different anti-CEACAM antibodies after *C. albicans* stimulation, respectively. Fig 2 highlights the differences in transcriptional responses elicited by either of the anti-CEACAM antibodies in presence of *C. albicans* compared to the IgG treatment in presence of *C. albicans*, i.e. the alterations obtained on top of the response induced by the *C. albicans* stimulation alone. Anti-CEACAM1 treatment in presence of *C. albicans* stimulation resulted in 19 upregulated DEGs compared to IgG treatment in presence of *C. albicans* stimulation (Fig 2, S3 Table), while anti-CEACAM3/*C. albicans* treatment led to no differential gene expression (Fig 2, S4 Table). Anti-CEACAM6/*C. albicans* treatment caused 75 DEGs compared to IgG/*C. albicans* treatment, with 65 upregulated and 10 downregulated genes (Fig 2, S5 Table). Systems biology analysis showed that 22 of the differentially regulated genes by anti-CEACAM6/*C. albicans* treatment were also regulated by *C. albicans*. Out of those 22 DEGs, 19 were synergistically co-regulated, and three were counter-regulated. GO enrichment analysis of DEGs altered by the anti-CEACAM6 antibody treatment in presence of *C. albicans* revealed that many are implicated in the regulation of major neutrophil functions, like cytokine production, neutrophil migration/chemotaxis, and cellular responses to pathogens, including TLR2-mediated responses (Fig 3, S9 Table). These data suggest that CEACAM6 ligation is likely to enhance important long-term responses initiated by neutrophils in response to *C. albicans*. Due to the fact that the anti-CEACAM1/*C. albicans* treatment shared the majority of regulated genes with the anti-CEACAM6/*C. albicans* treatment (17 of 19 DEGs, Fig 2), the GO enrichment analysis of CEACAM1-induced DEGs in presence of *C. albicans* resulted in less but similar results to those detected for the CEACAM6/*C. albicans*-treatment discussed above, like cell migration/chemotaxis and the regulation of responses to pathogens (S9 Table).

**Fig 2:**
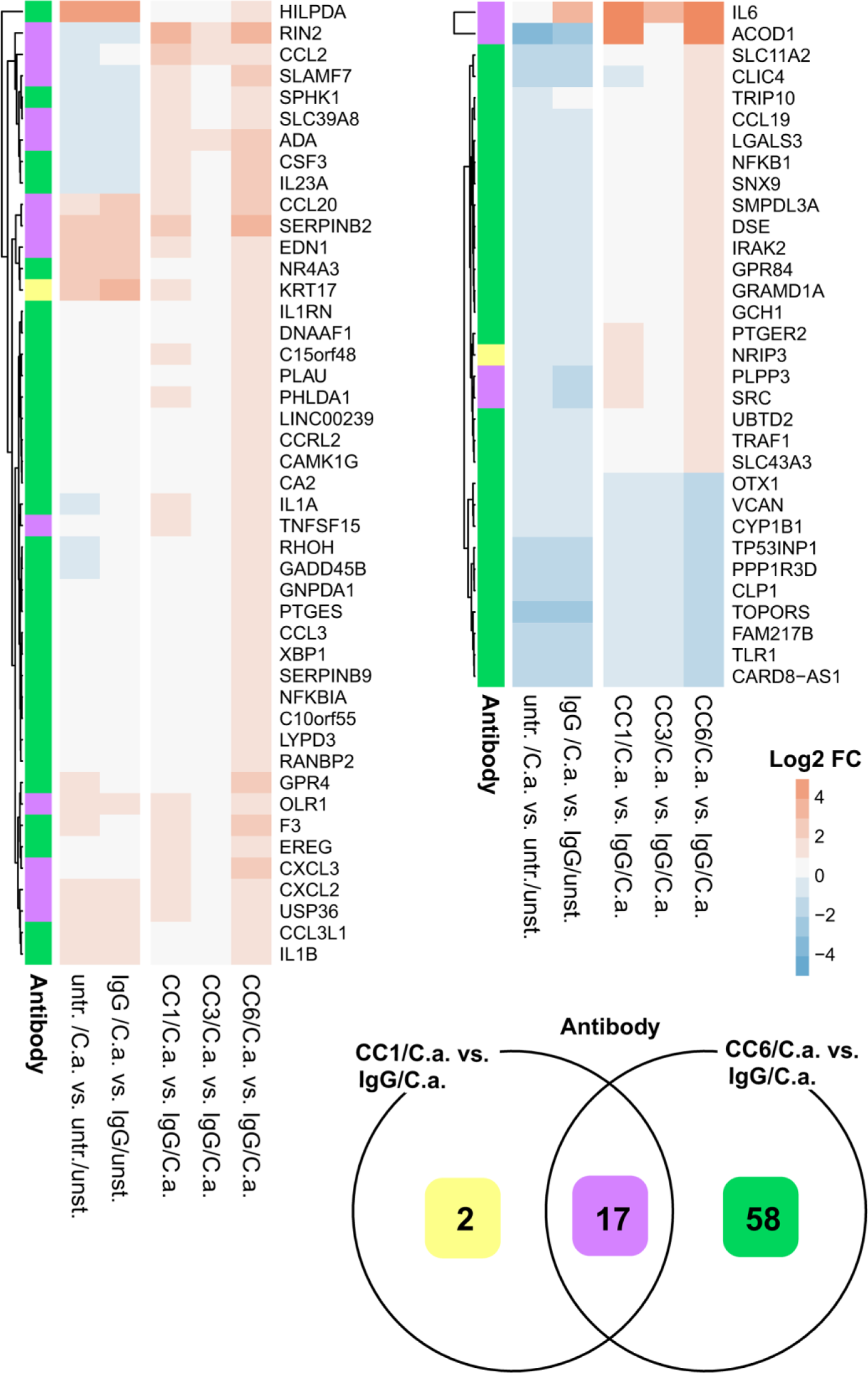
Altered Ca-induced neutrophil transcriptional response by priming with anti-CEACAM antibodies. In two independent experiments, human neutrophils were left untreated or were incubated with 10 µg/ml isotype control antibody (clone MOPC-21), or monoclonal antibodies B3-17 (CC1), 308/3-3 (CC3), and 1H7-4B (CC6), respectively, for 45 min and were consecutively incubated with or without live *C. albicans* yeast cells (MOI 1) for 2 h; mRNA was extracted and analyzed by sequencing. Differentially expressed genes (DEGs) from different comparisons are displayed in the heat map (untreated without C.a. vs. untreated with C.a., IgG-treated without C.a. vs. IgG-treated with C.a., IgG-treated with C.a. vs. CC1-treated with C.a., IgG-treated with C.a. vs. CC3-treated with C.a., and IgG-treated with C.a. vs. CC6-treated with C.a.). The latter three comparisons display the additional effect of the respective anti-CEACAM treatment on top of the transcriptional alterations induced by *C. albicans*; note that only significantly regulated genes with an adjusted p-value <0.05 and a fold-change of at least ±2 in one of these samples were included in the heat map. Each row was normalized (mean = 0) and scaled (standard deviation = 1). Which CEACAM treatment(s) altered the respective gene transcription is color-coded in the leftmost column of the heatmap and in the Venn diagram that also shows the total number of altered genes (yellow: unique for anti-CEACAM1 treatment in presence of *C. albicans*, green: unique for anti-CEACAM6 treatment in presence of *C. albicans*, purple: shared by anti-CEACAM1 and anti-CEACAM6 treatment in presence of *C. albicans*). Please refer to S1 to S5 Tables for details on DEGs as well as the complete lists of transcripts for all comparisons displayed in the heat map.

**Fig 3:**
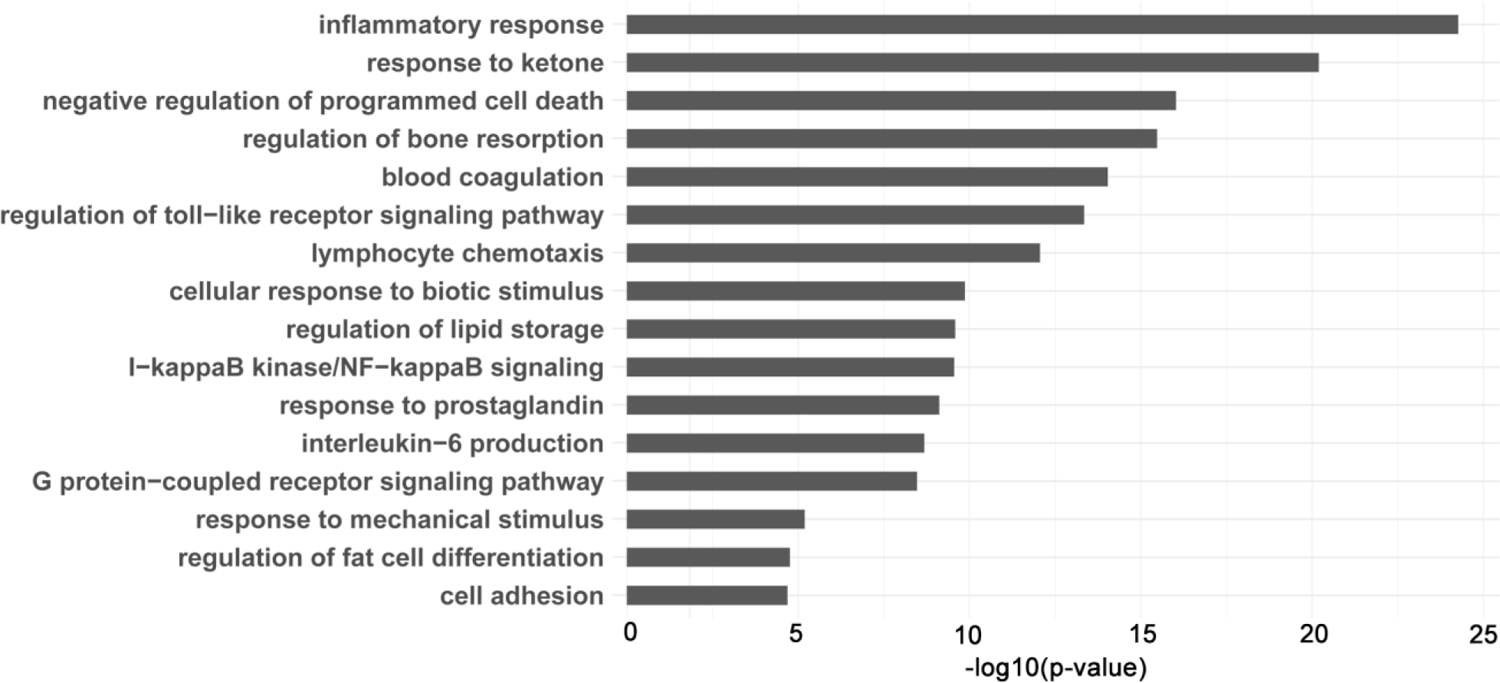
GO-enrichment analysis of CEACAM6-specific transcriptional responses of neutrophils in the presence of *C. albicans*. DEGs induced by CEACAM6 treatment in presence of *C. albicans* stimulation (CEACAM6-treatment in presence of *C. albicans* versus IgG-treatment in presence of *C. albicans;* data presented in Fig 2 and S5 Table) were analyzed for enriched GO terms and filtered for test values <0.01 in three different statistical tests (see Materials and Methods section for details). The graph displays compacted GO terms (consolidated according to semantic similarities) and their -log10(p-value). Please refer to S9 Table for the complete list of significantly enriched GO terms.

In absence of *C. albicans* stimulation, the anti-CEACAM1 antibody treatment resulted in 38 up-regulated and 1 down-regulated gene, and the anti-CEACAM6 antibody treatment in 151 up-regulated and 30 down-regulated genes (S5 Fig, S6 and S8 Tables). Again, the anti-CEACAM1 antibody treatment shared the majority of DEGs with anti-CEACAM6 antibody treatment (34 out of 39 DEGs; S5 Fig). Anti-CEACAM3 treatment shared its four upregulated genes with both, anti-CEACAM1 and anti-CEACAM6 treatments (S5 Fig, S7 Table). The GO enrichment analysis of differentially regulated genes by anti-CEACAM1 and anti-CEACAM6 antibody treatment in absence of *C. albicans,* respectively, revealed that also in absence of the *C. albicans* stimulation both antibodies regulated many cellular functions relevant for neutrophil immune responses, including migration/chemotaxis and TLR-dependent responses (S9 Table). Of course, the larger number of DEGs found after the anti-CEACAM6 antibody treatment also resulted in more enriched GO terms, including the production and secretion of cytokines and the regulation of apoptosis (S9 Table). Thus, priming via CEACAM1 and CEACAM6 ligation with B3-17 and 1H7-4B antibodies, respectively, is also likely to affect neutrophil functions in the absence of pathogens.

### Integrated network analysis and dynamic models predict the modulation of *C. albicans*-induced apoptosis and IL-1β production by anti-CEACAM6 antibody treatment

The gene expression data sets further allowed a systems biological analysis of the cellular functions and the signaling events involved. Since anti-CEACAM6 treatment had the most profound influence on the neutrophil transcriptional response in the presence of *C. albicans* stimulation, we further analyzed the cellular functions affected in dynamic models, and signaling pathways likely leading to these alterations in the cellular functions (Fig 4, S6 Fig, S10 and S11 Tables, described below). We considered signaling cascades triggered from the CEACAM6 ligation via direct protein-protein interactions as well as downstream pathways and functions induced subsequently according to the CEACAM6-specific transcriptional response (for detailed procedures refer to the Materials and Methods section).

**Fig 4:**
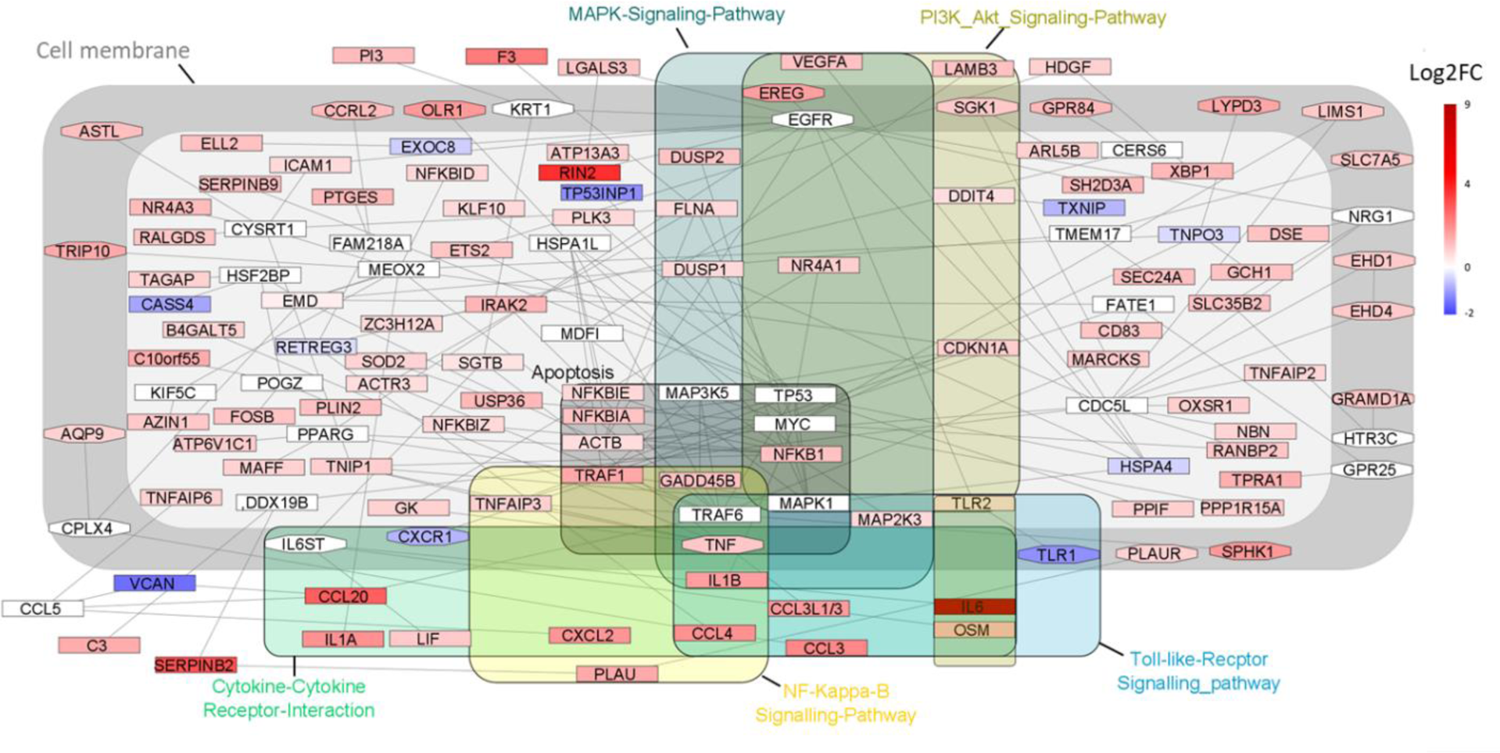
KEGG pathway enrichment analysis based on integrated network analysis of EACAM6-regulated genes in presence of *C. albicans*. Based on gene expression rofiles, the optimal CEACAM6-responsive network module in presence of *C. albicans* as identified. Subsequent KEGG pathway enrichment analysis detected key signaling athways of anti-CEACAM6 antibody-treated neutrophils stimulated with *C. albicans*. athway associated genes of selected pathways are highlighted and framed. Log2FC: log2 ld change. Main subcellular locations of gene products are indicated: intracellular, light rey area; membrane-associated, dark grey area; extracellular, outside grey areas. Please fer to S10 Table for a complete list of enriched pathways.

For the anti-CEACAM6 antibody treatment in the presence of *C. albicans* stimulation, the dynamical network analysis showed that the regulation of cytokine production and the regulation of apoptosis were among the significantly enriched cellular functions according to the Gene Ontology (GO) data base (S6 Fig, S11 Table). Also, the regulation of cell migration, and other basic functions important for the execution of neutrophil responses, including the regulation of TLR signaling, were enriched. When we performed a similar dynamical network analysis of the transcriptional data from the anti-CEACAM1-treated neutrophils in the presence of *C. albicans*, some similar immune-function-related GO categories were enriched, like cell migration and the regulation of cytokine production (S7 Fig). Interestingly, no apoptosis-related categories were enriched in the CEACAM1 network analysis.

Moreover, we performed an integrated network analysis for the anti-CEACAM6 treatment in presence of *C. albicans.* This yielded an optimally responsive network module of 136 genes with 174 interactions comprising distinguished key players annotated to important cellular functions including apoptosis and cytokine production. This is reflected by the highly significant enrichment of KEGG pathways related to these functions identified by the integrated network analysis (Fig 4, S10 Table), and is also in accordance with the GO-based analysis described above. The KEGG pathways overlapped to some degree, since many key signaling molecules are shared by different pathways. Central to the regulation of apoptosis and cytokine production are molecules of the NFκB complex and associated signaling partners, as illustrated in Fig 4. Moreover, dynamical simulations based on the transcriptomic data predicted the activation of caspase-8 (CASP8) and CASP1, two molecules recognized as central switches for apoptosis (41, 42), by anti-CEACAM6 treatment in presence of *C. albicans* stimulation, but not by CEACAM1/*C. albicans* treatment (S8 Fig). Interestingly, significantly enriched KEGG pathways of the anti-CEACAM6 treatment in presence of *C. albicans* stimulation also included the regulation of TLR2 and TLR4 signaling (Fig 4, S10 Table). So far, only for CEACAM1 and CEACAM3 experimental evidence is available for the regulation of these important pattern recognition receptors (8-10, 24, 43), which can recognize both, bacterial and fungal pathogens (44).

### Verification of the biological significance of the predicted increase in *C. albicans*-induced neutrophil apoptosis by anti-CEACAM6 ligation

Taken together, the predictions by the cell function-centered network analysis (S6 Fig) are well in agreement with the projections of the pathway-centered integrated network analysis (Fig 4) and the dynamical simulations (S8 Fig). The different bioinformatic approaches identified programmed cell death/apoptosis as one of the cellular functions most dominantly affected by CEACAM6 treatment in presence of *C. albicans* stimulation. *In vitro* experiments completely supported these predictions (Fig 5). CEACAM6 ligation with the specific monoclonal antibody 1H7-4B enhanced the *C. albicans*-induced apoptosis of human neutrophils, while none of the antibody treatments targeting CEACAM1 or CEACAM3 altered the neutrophil apoptosis (Fig 5A, B). Moreover, neither the anti-CEACAM6 antibody nor the anti-CEACAM1 and anti-CEACAM3 antibodies affected the spontaneous apoptosis (without *C. albicans* stimulation) of the neutrophils after 4 h and after 24 h, respectively (Fig 5C, D). Thus, the CEACAM6 ligation by 1H7-4B specifically enhanced the *C. albicans*-induced apoptosis.

**Fig 5:**
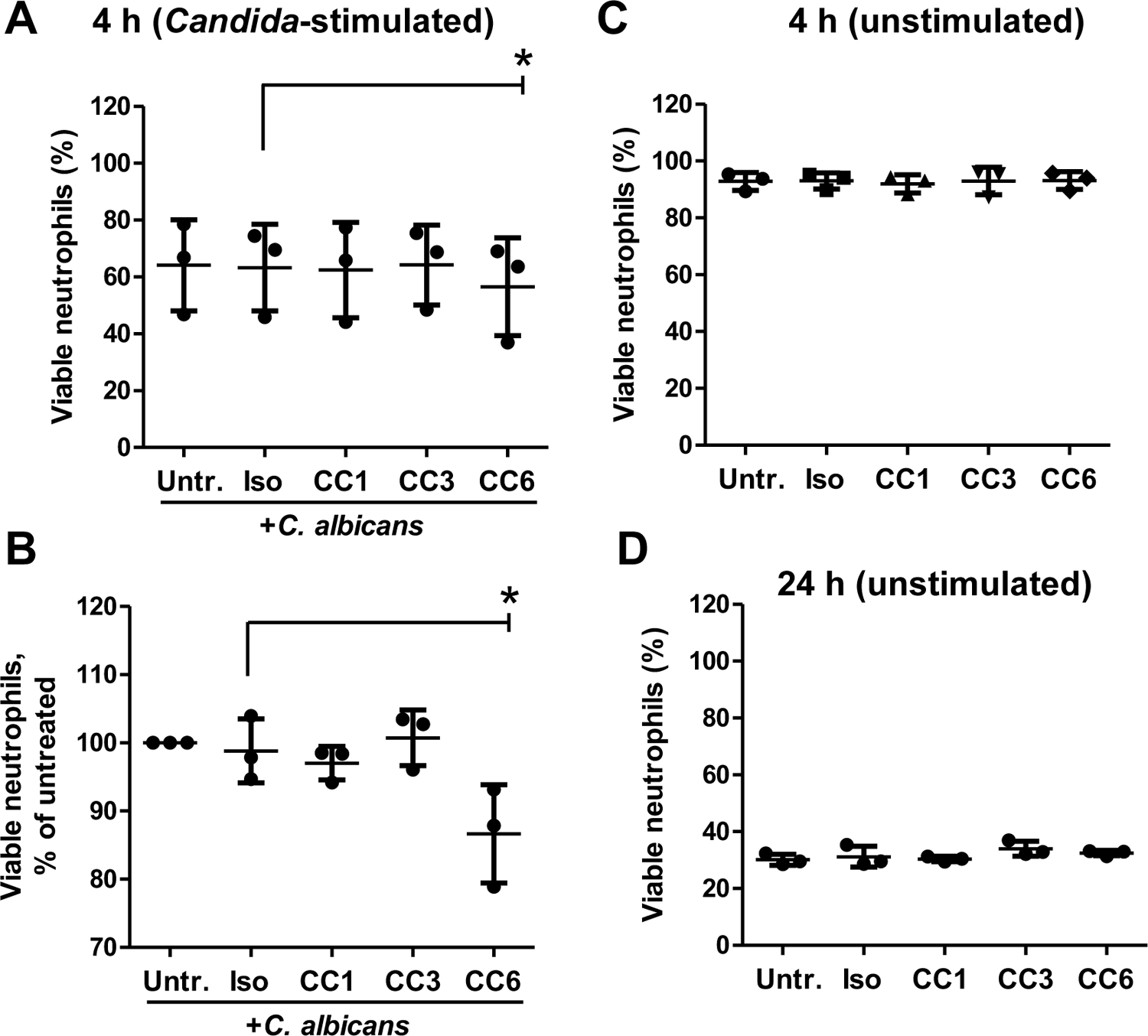
Verification of the altered apoptosis of human neutrophils after stimulation with *C. albicans* by anti-CEACAM6 antibody treatment predicted by different network analyses. Human neutrophils were left untreated or were incubated with the following antibodies: isotype control (Iso), B3-17 (CC1), 308/3-3 (CC3) or 1H7-4B (CC6) for 45 min. Afterwards, cells were incubated with or without live *C. albicans* yeast cells (MOI 1) for 4 h (A, B, C) or for 24 h (D). Viability of human neutrophils was determined by Annexin V and propidium iodide staining with subsequent flow cytometric analysis in three independent experiments. (A, C, D) graphs display the percentage of viable neutrophils (% of total neutrophils). (B) The graph displays the same samples shown in (A) as percentage of viable neutrophils compared to untreated cells in each experiment (viable untreated cells = 100%) to highlight the donor-independent relative effect of CEACAM6 treatment on *C. albicans*-induced apoptosis. Statistical analysis was performed by Repeated Measures ANOVA and Bonferroni post-test, *p<0.05.

### Verification of the biological significance of predicted alterations of neutrophil cytokine responses to *C. albicans* by anti-CEACAM1/3/6 ligation

For the predicted cytokine production, a more complex situation presented itself in the bioinformatic analyses as well as by a simple look at the transcriptional data. Transcription of the interleukin-1β gene (*IL1B)* was moderately but significantly induced by *C. albicans* stimulation alone of human neutrophils after 2 h (Fig6 A). It was further moderately enhanced by anti-CEACAM6 treatment in the presence of *C. albicans* stimulation, but also by anti-CEACAM1 treatment and anti-CEACAM6 treatment, respectively, in the absence of *C. albicans*, while anti-CEACAM3 treatment had no significant impact on the *IL1B* gene expression under any condition (Fig6 A). Network analyses and GO/KEGG enrichment all predicted a general activation of NFκB-dependent signaling and cytokine production induced by anti-CEACAM6 treatment in absence and in presence of *C. albicans* stimulation (Fig 4, S6 Fig, S9 Table). Dynamical simulations and GO enrichment even specifically predicted an enhanced IL-1β production by anti-CEACAM6 treatment (S8 Fig, S11 Table). In contrast, GO enrichment analysis for anti-CEACAM1-treatments in absence or presence of *C. albicans* stimulation did not show any significant alterations with respect to NFκB signaling and cytokine production in general or IL-1β production in particular (S9 Table). Also, the dynamic simulation (S8 Fig) predicted no activation of *IL1B* in presence of *C. albicans* stimulation. However, network analysis predicted a general regulation of cytokine production by the anti-CEACAM1 treatment in presence of *C. albicans* stimulation (S7 Fig). We therefore analyzed the secretion of the pro-inflammatory cytokine interleukin-1β (IL-1β) in detail in long-term experiments.

When we analyzed the secretion of IL-1β protein into the cell culture supernatants after 21 h (Fig 6B, C), we found that *C. albicans* stimulation indeed increased the total release of IL-1β from 5.7 ±1.5 pg/ml (“Untreated”, Fig 6B) to 40.0 ±12.4 pg/ml (“Untreated+*C. albicans*”, Fig 6C). IgG isotype treatment had no effect on the IL-1β release (Fig 6B, C). In accordance with the dynamic simulations (S8 Fig) and the GO enrichment (S9 Table), and in contrast to the CEACAM1 treatment-evoked increase in the *IL1B* transcripts (Fig 6A), CEACAM1 ligation did not result in detectable differences in the amount of secreted IL-1β compared to isotype-treated neutrophils neither in the presence nor absence of *C. albicans* stimulation (Fig 6B, C). Interestingly, CEACAM3 treatment led to a significant increase in IL-1β secretion with or without *C. albicans* stimulation (Fig 6B, C), despite the unaltered *IL1B* transcription levels after 2 h (Fig 6A). Concordant with the transcriptional response (Fig 6 A) and the bioinformatic predictions (Fig 4, S10 Table), CEACAM6 treatment resulted in a significant increase in IL-1β secretion in the presence of *C. albicans* stimulation at 21 h (Fig 6C). In the absence of the fungal pathogen, IL-1β levels were moderately but not significantly enhanced by CEACAM6 ligation at 21 h (Fig 6B).

**Fig 6:**
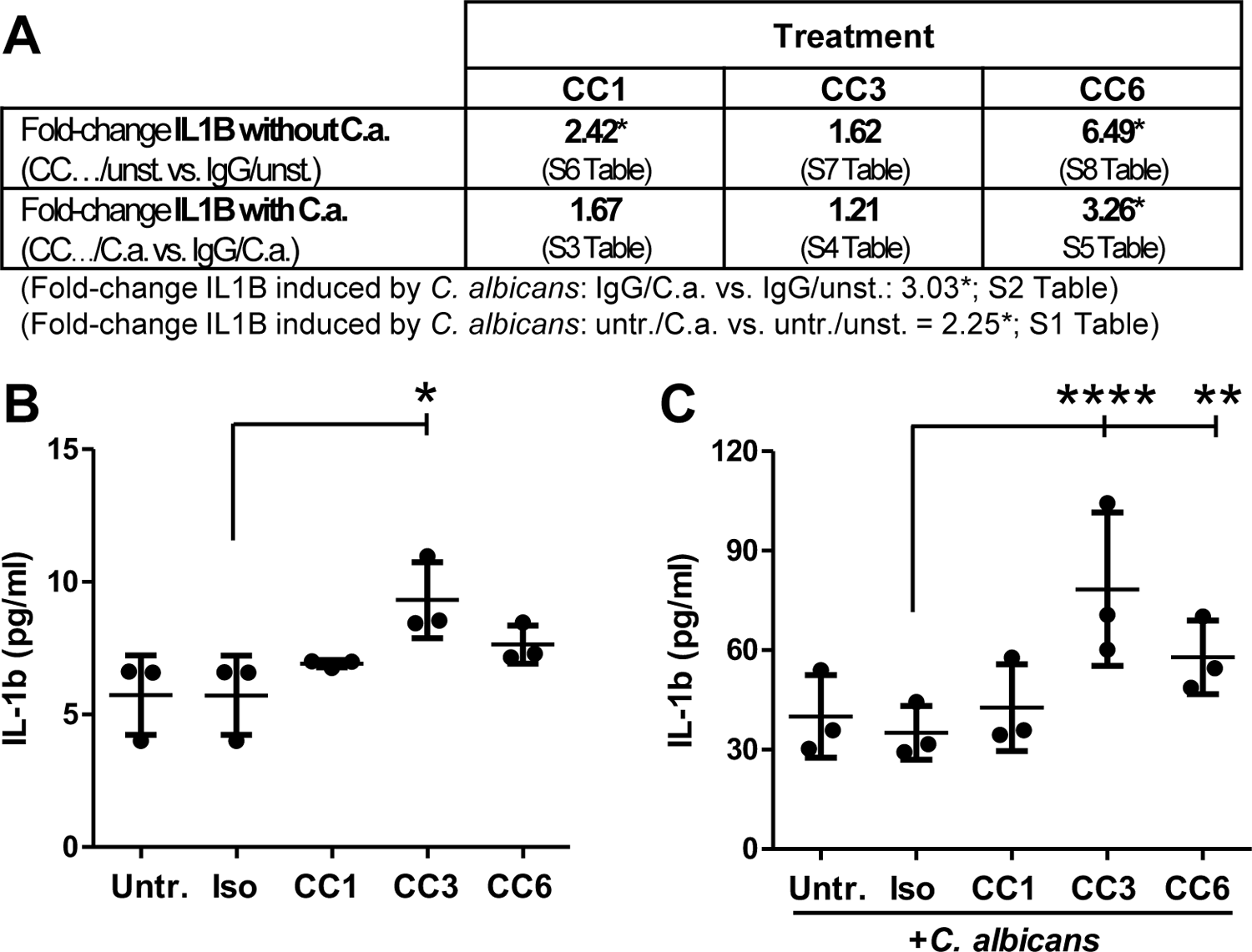
*C. albicans*-induced *IL1B* transcription after 2 h and IL-1β secretion after 21 h is regulated differentially by anti-CEACAM1, anti-CEACAM3, and anti-CEACAM6 treatment. Human neutrophils were left untreated or were incubated with the following antibodies: isotype control (Iso), B3-17 (CC1), 308/3-3 (CC3) or 1H7-4B (CC6) for 45 min. Afterwards, cells were incubated with or without live *C. albicans* yeast cells (MOI 1) for 2 h (A) or 21 h (B, C). (A) In two independent experiments, mRNA was extracted after 2 h and analyzed by sequencing (data from Fig 2 and S2 to S8 Tables). Fold-changes of the *IL1B* gene expression compared to isotype treatment in absence or presence of *C. albicans* stimulation, respectively, are shown. Fold-changes with an adjusted p-value<0.05 are marked with an asterisk. (B, C) Supernatants were collected after 21 h and analyzed for IL-1β concentrations in three independent experiments (sensitivity: 4 pg/ml). Statistical analysis was performed by Repeated Measures ANOVA and Bonferroni post-test; two samples below the detection range were set to the detection limit for statistical analysis.

The CEACAM3 ligation-induced IL-1β production, despite the lack of a detectable increase in the transcriptional activity after 2 h, led us to reinvestigate our network analysis. The sub-network analysis revealed a possible interaction between *IL6* and the subsequent induction of *IL1B* (Fig 7A), and *IL6* was among the four genes upregulated by anti-CEACAM3 treatment (S5 Fig, Fig 7B). Importantly, the regulation goes both ways, i. e. IL-1β is not only able to induce IL-6, but it can, *vice versa*, also be induced by IL-6. However, in contrast to the well-described “direct” induction of IL-6 via IL-1β, there were at least 4 nodes between *IL6* and *IL1B* (Fig 7A). *C. albicans* stimulation alone did not alter the *IL-6* transcription significantly, but it was significantly upregulated by all three anti-CEACAM antibodies in the absence of *C. albicans*, respectively (Fig 7B). However, in the presence of *C. albicans* only anti-CEACAM1 and anti-CEACAM6 treatment reached significance for the increase in *IL-6* transcription (Fig 7B). This was likely due to the very low *IL-6* copy numbers detected by sequencing: in untreated and isotype-treated neutrophils, both, in the presence and the absence of *C. albicans* stimulation, respectively, the normalized counts of the *IL6* transcripts in all samples were between 0 and 3. Normalized *IL-6* counts in all samples of the anti-CEACAM treatment groups with or without *C. albicans* stimulation were in the range between 24 and 1,447, resulting in the high variance for the *IL6* transcripts. By comparison, the lowest normalized count detected for *IL1B* transcripts was 18,023 in an untreated sample without *C. albicans* stimulation. Thus, very few transcripts serve neutrophils for the *de novo* synthesis of the pro-inflammatory cytokine IL-6.

**Fig 7:**
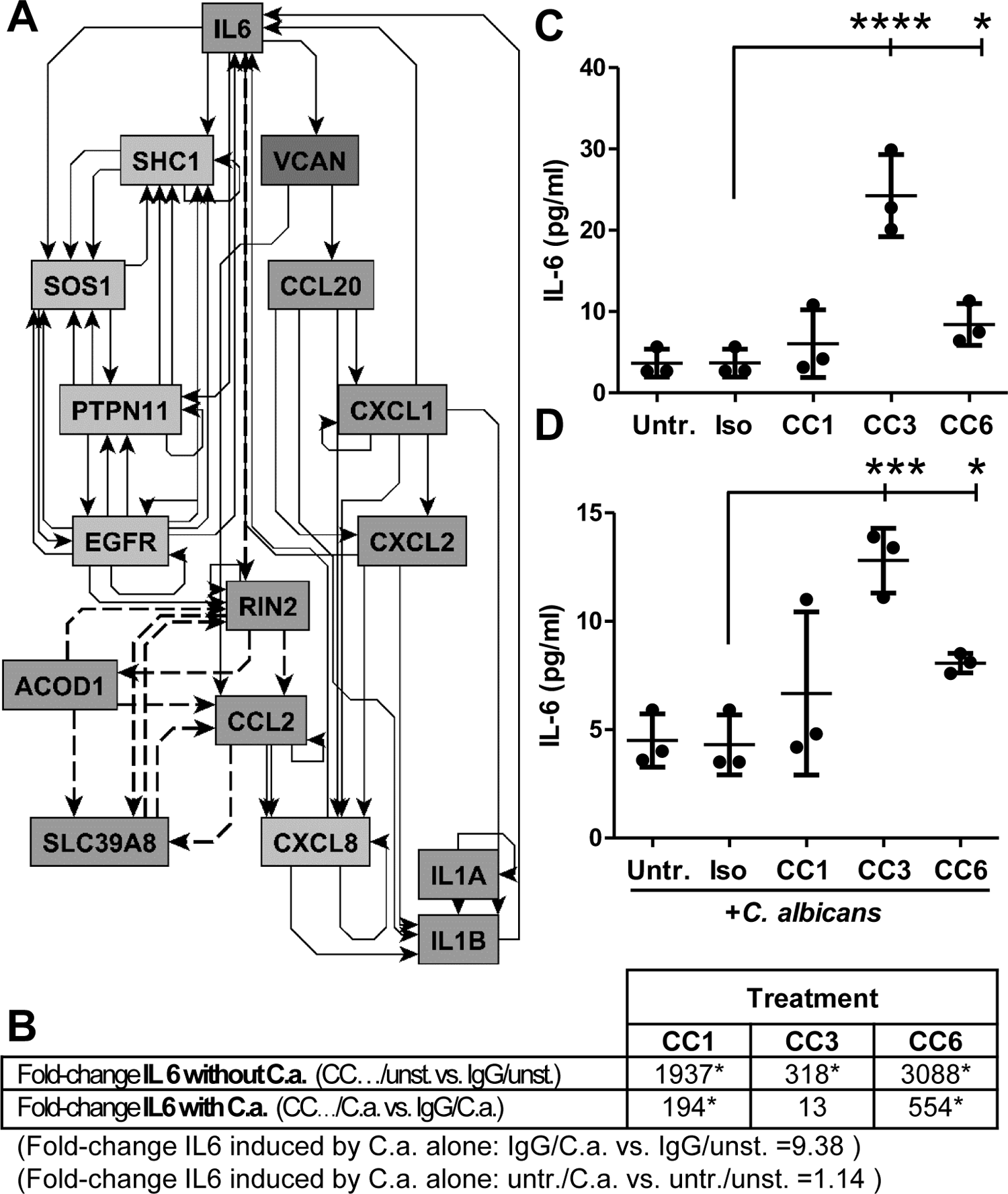
IL-6 secretion after 21 h is induced by ligation of CEACAM3 and CEACAM6 but not of CEACAM1. (A) Sub-network of *IL1B* induction by IL6 signaling. The network shown is a subgraph from the network displayed in Figure S7 (data from Figure 2). All paths between IL6 and *IL1B* up to 7 nodes long were extracted; dashed lines: inferred interactions from our network analysis, solid lines: interactions obtained from data bases. (B-D) Human neutrophils were left untreated or were incubated with the following antibodies: isotype control (Iso), B3-17 (CC1), 308/3-3 (CC3) or 1H7-4B (CC6) for 45 min. Afterwards, cells were incubated with or without live *C. albicans* yeast cells (MOI 1) for 2 h (B) or 21 h (C, D). (B) In two independent experiments, mRNA was extracted after 2 h and analyzed by sequencing (data from Figure 2). Fold-changes of the *IL6* gene expression compared to isotype treatment in absence or presence of *C. albicans* stimulation, respectively, are shown. Fold-changes with an adjusted p-value<0.05 are marked with an asterisk. (C, D) Supernatants were collected after 21 h in three independent experiments and analyzed for IL-6 concentrations (sensitivity: 2 pg/ml). Statistical analysis was performed by Repeated Measure ANOVA and Bonferroni post-test.

To analyze whether these few transcripts were translated into detectable protein levels secreted after contact with *C. albicans*, we measured IL-6 levels in neutrophil supernatants after 21 h of infection (Fig 7C, D). The lack of transcriptional *IL6* expression and induction by *C. albicans* was verified *in vitro* via ELISA, where 3.7±1.7 pg/ml IL-6 were produced by untreated neutrophils in the absence, and 4.5±1.2 pg/ml IL-6 in the presence of *C. albicans* (Fig 7C and D, respectively; 10^7^ neutrophils/ml). IgG treatment did not change IL-6 levels of unstimulated or *C. albicans*-stimulated neutrophils. CEACAM3 treatment significantly enhanced the IL-6 secretion in the absence and the presence of *C. albicans*, respectively (Fig 7C, D). CEACAM6 treatment augmented the IL-6 release with and without *C. albicans* stimulations slightly but significantly (Fig 7C, D). CEACAM1 treatment only resulted in minor, non-significant increases in IL-6 secretion in neutrophils from one preparation, both, in presence and in absence of *C. albicans* stimulation (Fig 7C, D), despite a transcriptional induction at 2 h (Fig 7A).

## Discussion

Here, we show for the first time that the ligation of CEACAM receptors expressed on human neutrophils by specific antibodies influence the human neutrophil response to *C. albicans*. In particular, the ligation of CEACAM6 by the 1H7-4B antibody in the presence of *C. albicans* stimulation enhanced the early release of CXCL8 and altered the neutrophil transcriptional responses. Different bioinformatic approaches (transcriptome analysis, network analysis and dynamical modelling of the involved response) then led to the identification of long-term neutrophil responses modulated by the CEACAM6-treatment in *C. albicans*-stimulated neutrophils, i.e. *C. albicans*-induced apoptosis and IL-6/IL-1β *de novo* synthesis. Ligation of CEACAM1 by the B3-17 antibody enhanced the early CXCL8 release, and CEACAM3 ligation by the 308/3-3 antibody increased the late IL-6 and IL-1β production.

It was surprising to find that the secretion of CXCL8 and the global transcriptional response after 2 h of *C. albicans*-stimulation were enhanced by the antibody-mediated ligation of CEACAM1 and CEACAM6, but not of CEACAM3. On neutrophils, CEACAM3 is recognized as the central CEACAM family receptor leading to neutrophil activation by CEACAM-binding bacteria, and the subsequent phagocytosis and bacterial killing due to an ITAM in its cytoplasmic tail (23-25, 28, 45-49). Despite the dominance of CEACAM3 in the human neutrophil responses to pathogenic bacteria, both, CEACAM1 and CEACAM6, can also mediate binding and phagocytosis of bacterial pathogens in neutrophils and in other cell types (28, 49, 50). Also, CEACAM receptor ligation by various antibodies results in the activation of neutrophils resulting in an enhanced binding to endothelial cells (33, 34). On the other hand, CEACAM1 negatively regulates the TLR-4 dependent IL-1β production in LPS activated neutrophils (10). In epithelial cells, CEACAM1 induces either inhibitory or activating effects on their immune responses; e.g. in lung epithelial cells, CEACAM1 dampens TLR2-mediated responses (8, 18), while it is critical for the pathogen-induced CXCL8 release in gastric and intestinal cell lines (12, 17). Further work will be necessary to find out whether the CEACAM1- and the CEACAM6-mediated effects described in the present study also depended on the regulation of TLR family receptors as implicated by the different bioinformatic analyses used.

CEACAM1 can inhibit the function of another essential receptor on mouse neutrophils, granulocyte-stimulating growth factor receptor, resulting in a reduced granulopoiesis (32). And on mouse monocytes, CEACAM1 can negatively regulate IL-6 production (51). These dampening effects of CEACAM1 on the mouse myeloid responses enhanced the susceptibility of mice to bacterial pathogens and LPS challenge *in vivo* (32, 51). In accordance with our findings in epithelial cells (12, 17), CEACAM1 ligation by B3-17 antibody enhanced the immediate pro-inflammatory CXCL8 secretion by neutrophils in response to *C. albicans*. Interestingly, the long-term response of the neutrophils seemed not affected by CEACAM1 ligation, since the two important pro-inflammatory cytokines IL-6 and IL-1β were not enhanced in the CEACAM1 treated neutrophils in the presence and the absence of *C. albicans* stimulation, respectively, despite the increase in *IL1B* and *IL6* transcripts. The precursor of IL-1β is biologically inactive and requires proteolytic cleavage into biologically active mature cytokines, controlled by the inflammasome-mediated caspase-1 activation (52). In fact, in mouse neutrophils, CEACAM1 negatively regulates the LPS-induced IL-1β production by blocking caspase-1 activation (10). However, our experiments showed no alteration of IL-1β levels at all after B3-17 treatment in the presence and the absence of *C. albicans* stimulation.

It should also be noted that not all specific antibodies used in this study evoked the same effects or affected neutrophil functionality to variable magnitudes when CEACAM1, CEACAM3 or CEACAM6 were ligated. Different antibodies to CEACAM1 can either enhance or block CEACAM1-mediated adhesion in trans and thereby can modulate the dimerization status of CEACAM1 or intracellular binding of SHP1 and SHP2 phosphatases and Src-like kinases to the CEACAM1-long cytoplasmic domain (53). Furthermore, it is well established that CEACAM1 ligation on epithelial cells can transduce inhibitory or activating cell signaling events dependent on the cellular status or context of the same cell (54). Taken together, our data disclose a role for CEACAM1 ligation in increasing the immediate pro-inflammatory neutrophil response to C*. albicans*, in particular the enhanced CXCL8 secretion.

CEACAM3 played a very different role in the neutrophil responses to *C. albicans*. While there were no alterations upon CEACAM3 ligation in the immediate neutrophil responses (CXCL8 release, fungal killing), and no genes were significantly upregulated after 2 h of *C. albicans* stimulation, CEACAM3 treatment resulted in the highest concentrations of all treatment groups for IL-1β and IL-6 after 21 h incubation. Thereby, CEACAM3 ligation showed the highest potential of increasing local inflammation with all its cytotoxic side effects (2, 4). Taken together, the role of CEACAM3 in the regulation of neutrophil responses to the fungal pathogen studied here, and to various bacterial pathogens discussed earlier, differs widely: CEACAM3 has a major activating effect on the immediate response to the bacterial pathogens (23-25, 28, 45-49), but regulates the long-term cytokine release enhancing the overall inflammatory response in *C. albicans* infection by the subsequent attraction of additional immune cells.

Increased CEACAM6 expression levels on neutrophils are correlated with asthma severity (55), and the authors of that study propose the presence of an altered neutrophil phenotype in presence of increased CEACAM6 expression. As stressed before, CEACAM6 ligation by antibodies can activate neutrophils and induce enhanced binding to endothelial cells (33, 34). We believe this is the first study showing that anti-CEACAM6 ligation enhanced neutrophil cytokine production. Our experiments also likely underestimated the levels of IL-1β and IL-6 produced per neutrophil upon anti-CEACAM6 treatment with the 1H7-4B antibody during *C albicans* infection, since the same treatment also led to an increase in apoptosis, thereby reducing the numbers of neutrophils contributing to the cytokine levels. An increase of CEACAM6 expression on epithelial cells and leukocytes is induced by various pathogens and pro-inflammatory cytokines (12, 56–59) and is thus generally associated with inflammatory events. CEACAM6 interaction on ileal mucosa with adherent-invasive *E. coli* (AIEC), bacterial pathogens associated with Crohn disease (CD), results in an enhanced inflammation *in vivo*, both, in CD patients and in a CD mouse model using CEABAC10 transgenic mice (59, 60). Further, CEACAM6-AIEC interaction also enhances cytokine production in mucosal cells *in vitro* (56, 59).

For the anti-CEACAM6 ligation, all three bioinformatic approaches consistently predicted an influence on the *C. albicans*-induced apoptosis. In contrast, no such prediction arose for the anti-CEACAM1 ligation. This was rather surprising, since our earlier publication shows that in rat granulocytes, CEACAM1 delays the spontaneous and the FAS ligand-mediated apoptosis (20), while on lung epithelial and colorectal cancer cells, CEACAM1 supports apoptosis (61, 62). In the experimental set-up used here, only CEACAM6 ligation by 1H7-4B in presence of *Candia*-stimulation lead to an enhanced apoptosis. All other treatments did not alter survival of the neutrophils compared to untreated neutrophils in presence or absence of *C albicans* stimulation, respectively. So far, most studies show that CEACAM6 mostly decreases apoptosis and anoikis (apoptosis induced by the lack of adhesion) in various cancer cells and in primary epithelial cells (63–69). Interestingly, one study on acute lymphoblastic leukemia also demonstrates an increase in apoptosis via antibody-mediated ligation of CEACAM6 (70).

Neutrophils co-express not only CEACAM1, CEACAM3 and CEACAM6, but also CEACAM8. Of these, the former three are likely co-stimulated by *C albicans* and able to modulate the functions of other members of the CEACAM family (34). Therefore, further experiments will be necessary to determine the exact roles of each single CEACAM family member during neutrophil responses to *C albicans*, and whether one CEACAM receptor appears dominant in the response to the fungal pathogen. Since neutrophils are short-lived and therefore are not useful for the transfection with siRNA, ligation with antibodies remains the best accessible tool. Any CEACAM-monospecific or even cross-reacting antibody (mimicking co-ligation by the pathogen) has to be evaluated on its own and in the specific setting of the experiment, since targeting different epitopes on the same CEACAM receptor can either enhance or inhibit the receptor functions (53), and one antibody can elicit inhibitory or activating cell signaling events dependent on the cellular condition of the same cell (54). While neutrophils from transgenic mice expressing one or more human CEACAMs proved valuable for the determination of the role of CEACAM receptors in the neutrophil response to bacterial and viral pathogens (22, 28, 31, 46), our data obtained with *C albicans* stimulated neutrophils from human CEACAM1-transgenic mice suggest that mouse neutrophils do not respond to the fungal pathogen in a CEACAM1-dependent manner. Since mice are inherently naïve for *C albicans* infection, it is possible that some (co-)receptor or signaling protein important for the CEACAM-specific response to this human pathogen is lacking in the mouse neutrophils.

Taken together, the combination of transcriptome analysis, systems biological modelling, and cytokine measurements in the present study revealed that antibody-mediated ligation of CEACAM1, CEACAM3, and CEACAM6 could elicit specific regulations of neutrophil responses, evoked by the fungal pathogen *C albicans.* CEACAM1 ligation had an early pro-inflammatory effect on human neutrophils during *C albicans* infection (CXCL8 release). CEACAM3 ligation had no early effects during *C albicans* infection, but acted strongly pro-inflammatory in long-term experiments (IL-1β and IL-6 production). CEACAM6 ligation consistently displayed an activating activity in neutrophils in early (CXCL8 release, transcriptome) and long-term (IL-6 and IL-1β release) responses. Surprisingly, CEACAM6 ligation also resulted in an enhanced *C albicans*-induced apoptosis.

In summary, the results of our study demonstrate for the first time that the interaction of *C. albicans* with CEACAM1, −3, and −6 has important and differential immunomodulatory effects on the regulation of transcriptional responses, cytokine release, and cell death of human neutrophils. These results suggest that CEACAM receptors play an important role in *Candida albicans*-induced immune responses of human granulocytes as first line defenses against fungal pathogens.

## Materials and Methods

### Neutrophil isolation and treatments

Human peripheral blood was collected from healthy volunteers with written informed consent. This study was conducted according to the principles expressed in the Declaration of Helsinki. The blood donation protocol and use of blood for this study were approved by the institutional ethics committee of the University Hospital Jena (permission number 5070-02/17). Neutrophils were isolated from peripheral blood using 1-Step Polymorphs (GENTAUR GmbH, Germany). In brief, 20 ml 1-Step Polymorphs were overlaid with 20 ml blood and centrifuged at 500 × g for 35 min at room temperature with acceleration and deceleration set to “0”. The lower cell layer was collected, mixed with one volume of ice-cold 0,45% NaCl solution, and centrifuged at 4°C and 400 × g for 10 min. the pellet was suspended in 5 ml ice-cold ACK lysis buffer. The reaction was stopped by adding 45 ml HBSS (GIBCO, Thermo Fisher Scientific, Germany). Centrifugation for pelleting and a further washing step using HBSS were performed at 4°C and 250 × g for 10 min. Purity >96% was determined by flow cytometry. Cells numbers were adjusted to 1×10^7^ cells/ml in RPMI/10% FBS if not stated otherwise and experiments were performed immediately. Cells were either left untreated or treated with 10 µg/ml antibody (see below) for 45 min. Cells were then either left unstimulated or stimulated with *C. albicans* for 2 h at an MOI of 1 if not mentioned otherwise.

### Antibodies and recombinant CEACAM6-Fc

All antibodies used were monoclonal mouse IgG1: MOPC-21 (Hoelzel Diagnostika GmbH, Germany; isotype control), B3-17 (anti-human CEACAM1, Singer, Essen, Germany), C5-1X/8 (anti-human CEACAM1, Singer, Essen, Germany), CC1/3/5-Sab (anti-human CEACAM1/3/5, LeukoCom, Essen, Germany), 18/20 (anti-human CEACAM1/3/5, Singer, Essen, Germany), 308/3-3 (anti-human CEACAM3/5, LeukoCom, Essen, Germany), 1H7-4B (anti-human CEACAM6, LeukoCom, Essen, Germany), 13H10 (anti-human CEACAM6, Genovac, Freiburg, Germany). Specificity of all antibodies was verified by FACS analysis using CHO or HELA cells transfected with human CEACAM1, CEACAM3, CEACAM6, CEACAM5, CEACAM8, or empty vector (negative control). Note that none of the antibodies cross-reacted to CEACAM8, and that 18/20, CC1/3/5-Sab, and 308/3-3 cross-reacted to CEACAM5 that is not expressed on neutrophils (therefore we refer to 308/3-3 as “CEACAM3-specific” in the context of this publication). Cross-reactivities to CEACAM1, CEACAM3, and CEACAM6 are given in Fig 1. Recombinant CEACAM6 protein consisting of the CEACAM6 extracellular domain fused to the constant region of human IgG were produced in HEK-293 cells and purified via protein G columns (GE Healthcare, Munich, Germany) as described previously (71).

### Candida albicans culture

*C. albicans* strain SC5314 was grown as described (72, 73). For experiments, YPD liquid cultures were inoculated with a single colony from YPD agar plates and grown at 30°C and 180-200 rpm for 14-16 h. Yeast cells were harvested, washed twice and suspended in a desired volume of ice-cold PBS. Yeast cells were counted in a Neubauer Improved chamber.

### ELISA

1×10^7^ neutrophils/ml in RPMI/10% FBS were either left untreated or treated with 10 µg/ml antibody for 45 min. Cells were then either left unstimulated or stimulated with *C. albicans* for 2 h at an MOI of 1. Supernatants were harvested at the indicated time points and tested by ELISA for CXCL8 (BD Biosciences, Germany; sensitivity: 3 pg/ml), IL-6 (Abcam, Germany; sensitivity: 2 pg/ml) and/or IL-1β (BD Biosciences, Germany, sensitivity: 4 pg/ml) concentrations.

### Binding analysis and ROS determination

Neutrophils were stained with anti-CD11b-PerCP-Vio700 (REA592, Miltenyi Biotech, Germany; 5 µl for 4×10^6^ neutrophils in 400 µl medium) for 45 min. *C. albicans* cells were stained on ice with Rabbit anti-*C. albicans* IgG (BP1006, Acris Antibodies, 5 µl for 6×10^7^ *C. albicans* cells in 200 µl medium) for 30 min, followed by Goat anti-Rabbit-AlexaFluor633 (Invitrogen, Sweden; 2 µl in 400 µl medium) for 30 min. 2×10^5^ stained neutrophils in 200 µl RPMI/10% FBS were left untreated or treated with 10 µg/ml antibody and incubated for 45 min at 37°C, 5% CO_2_. Neutrophils were left unstimulated or were stimulated with 1×10^6^ *C. albicans* cells per well (MOI 5) and samples were incubated for 30 min at 37°C, 5% CO_2_. 10 µl of 0.5 µg/ml DHR (Sigma-Aldrich, Germany) working solution in 4% FBS/D-PBS (Gibco/Thermo Fisher Scientific, Germany) was added to each sample to reach a final concentration of 25 ng/ml, and samples were incubated at 37°C, 5% CO_2_ for 10 min. Samples were analyzed in an Attune flow cytometer (Invitrogen/ Thermo Fisher Scientific, Germany; BL1: DHR123/ROS, BL3: CD11b/neutrophils, RL1: C. albicans). For adjustments and compensation, single stainings were performed before the experiments. Relative fluorescence units of the DHR123/ROS signals were logarithmized for presentation.

### *C. albicans* killing assay

In a 96 well plate, 2×10^5^ neutrophils in 100 µl RPMI/10% FBS were left untreated or treated with 10 µg/ml antibody in duplicates and incubated for 45 min at 37°C, 5% CO_2_. Neutrophils were left unstimulated or were stimulated with 2×10^5^ *C. albicans* cells per well (MOI 1) and samples were incubated for 30 min at 37°C, 5% CO_2_. *C. albicans* solution for standards (input) was kept on ice for the incubation times. Just before the following procedures, *C. albicans* standards (sensitivity: 6,500 CFU) were transferred to the 96 well plate. Triton X-100 was added to all wells to reach a final concentration of 0,3% and incubated for 10 min at 37°C, 5% CO_2_, in order inactivate neutrophils. Viable *C. albicans* cells were quantified using the Colorimetric Cell Viability Kits III (XTT) from Promokine, Germany: 50 µl XTT reaction mixture was added per well and samples were incubated for further 3-4 h at 37°C, 5% CO_2_. Absorbance was measured using a TECAN M200 at 450 nm and 630 nm (background). For calculations of *C. albicans* CFUs, background and blanks were subtracted.

### Spontaneous and *C. albicans*-induced Apoptosis

2×10^5^ neutrophils in 200 µl RPMI/10% FBS were left untreated or treated with 10 µg/ml antibody in duplicates and incubated for 2 h at 37°C, 5% CO2. Neutrophils were left unstimulated (spontaneous apoptosis) or were stimulated with 2×10^5^ C. albicans cells per well (MOI 1; C. albicans-induced apoptosis) and samples were incubated for 4 h or 24 h (unstimulated only) at 37°C, 5% CO2. Cells were stained with the Annexin V Detection Kit APC (eBioscience, Germany) and analyzed in an Attune flow cytometer (Invitrogen/ Thermo Fisher Scientific, Germany; BL2: propidium iodide, RL1: Annexin V-APC). Cells negative for both stainings were considered viable.

### RNA sequencing

1.2×10^7^ neutrophils in 1.2 ml in RPMI/10% FBS were either left untreated or treated with 10 µg/ml antibody for 45 min. Cells were then either left unstimulated or stimulated with *C. albicans* for 2 h at an MOI of 1. RNA was isolated using the Innuprep RNA minikit 2.0 0 (Jena Analytik, Germany) with an additional DNAse digestion step (RNase-free DNase set; Qiagen, Germany). Quality and quantity of the total RNA samples were evaluated with the Tape Station 2200 (Agilent, USA) and the Nanodrop 1000 (Thermo Fisher Scientific, Germany), respectively. Stranded RNA libraries were then prepared from 1 μg total RNA using the TruSeq Stranded mRNA Prep Kit (Illumina, USA) following the manufacturer’s protocol. Multiplexing of the samples was achieved using IDT–TruSeq RNA UD Indexes (Illumina, USA). The libraries were then loaded on a S1 Flowcell using the Xp Workflow and subjected to 100 cycles paired-end sequencing on the Illumina NovaSeq 6000 System.

### Analysis of sequencing data

Our dataset is composed by 40 fastq files corresponding to reads one and two (paired-end sequencing) of the 20 samples of the project. Those 20 samples correspond to replicates one and two of the 10 different conditions. All these files can be found in the SRA repository (https://www.ncbi.nlm.nih.gov/sra/PRJNA681392). We used the galaxy-europe server (74) to map the fastq files and count the gene transcriptions; the following named functions are all included in the server and were used using the default parameters except where it is otherwise indicated. First, the quality of the fastq files was assessed using the function *FastQC* then the adapter sequences were trimmed with the help of the function Cutadapt. This result was mapped to the human genome hg38 using the *RNA Star* applied for pair-ended sequences, the “Length of the genomic sequence around annotated junctions” parameter was set to 50. From the RNA Star function, we obtained the bam files which were quality controlled and later used to look for gene counts with the *FeatureCount* function. We processed the reverse stranded bam files allowing for fragment counts but not multimapping. The minimum mapping quality per read was set to 10. Finally, we obtained the count files for each of the samples in the project.

### DEG fold-change analysis

We analyzed the count files in R, using the *Deseq2* (75) and the *edgeR* (76) packages. The first step was to calculate the RPKM (Reads Per Kilobase Million) values for each gene, and discard those genes with an RPKM value lower than three. The remaining genes were processed to look for differentially expressed genes (DEG’s). For this purpose, a *deseq* object was created with the design formula *C. albicans + antibody* meaning that the antibody treatments were being analyzed while controlling for the *C. albicans* stimulus. The results for different contrasts are presented in S1 to S8 tables.

### PCA

The principal component analysis (PCA) was made directly with the function embedded in the *Deseq2* package, using the top 500 genes but no significant difference was observed when using the full set of genes. The antibody treatment was marked as interest group.

### Gene Ontology analysis

The GO terms were calculated using the topGO package in R (77). Three statistical tests were applied to the search, two Fisher exact tests, using the classic and the weight algorithms and a Kolmogorov-Smirnov test using the classic algorithm. For more details in these calculations please see the R package documentation. A total of 1000 terms were extracted with their respective test values. The lists were then filtered to keep only those terms which had all test values lower than 0.01. The obtained sets of GO terms were further compacted by applying the ReviGO algorithm (78) which summarizes GO terms lists based on semantic similarities. The resumed list of GO terms is plotted in the barplot, and the values of the bars correspond to the weighted Fisher exact test.

### Heatmaps

The heatmaps present the normalized count values of the mentioned genes averaged over the two replicates of each sample. Normalization assumed normal distribution. The normalization was done using the *counts* function of the *Deseq2* package with the parameter normalization set to true. Furthermore, within each subpanel heatmap the values were scaled (to mean = 0 and standard deviation = 1) by row, in order to facilitate the comparison between conditions (these are no longer absolute values).

### Subnetwork and Module analysis with subsequent KEGG pathway enrichment analysis

The network of human protein-protein interactions has been established in the FungiWeb database (C.H. Luther, C.W. Remmele, T.Dandekar, T. Müller, and M. Dittrich, submitted for publication) (79). This network contains protein-protein interactions, which have been collected and curated from experimental data sets from the International Molecular Exchange (IMEx) consortium (80) via PSIQUIC queries (81) and have been filtered and curated to focus on experimental interactions only. Furthermore, interactions have been scored analogously to the MINT-Database score (81), which reflects the quality and quantity of experimental information supporting each interaction. For integrated network analysis only high and highest quality interactions (score cutoff ≥0.75) have been used and a network comprising the largest connected component has been derived with a total of 17.754 genes and 237.846 interactions. The integrated network analysis aims to identify the maximally responsive regions (modules) after stimulation within the large interactome network, according to the integrated gene expression profiles. Thus, for the network analysis, RNA-Seq data was first preprocessed by an in-house pipeline, and differential gene expression was analyzed using a generalized linear model as implemented in DESeq2 (75), contrasting the different CEACAM-antibody treatments to the isotype antibody control. To obtain gene scores for the identification of responsive subnetworks a BUM (Beta Uniform Mixture) model has been fitted to the distribution of P-values using the routines in the BioNet package (82) choosing a stringent FDR (False discovery Rate) threshold of 10^-19^ to focus only on the maximally responsive region. Based on the scored network optimally responsive modules have been identified using an exact approach, which, albeit computationally demanding, is capable (mathematically proven) to identify provably optimal modules (83). Visualization of the resulting subnetworks has been performed using Cytoscape (84). Enrichment analysis of the network modules has been performed with g:Profiler (85) against the background set of the genes in the interaction network, using default settings and focusing on KEGG (86) pathways only. The information for cell membrane and secreted localization were obtained from UniProt (87). All statistical analyses have been performed with the computational statistics framework R (version 3.6.3).

### Systems biological modeling and network analysis with subsequent GO term enrichment analysis

We created protein signaling networks for CC6 (S6 Fig) and CC1 (S7 Fig). These networks combined the gene regulatory data from the RNA-sequencing analysis with data base knowledge of protein-protein interactions. First, the relevant genes were selected as those which reached an adjusted p-value lower than 0.05 and resulted in an expression of at least two-fold (red and blue nodes in S6 and S7 Figs. In addition to these genes, we assembled a set of known interacting proteins of the CEACAM receptors (gray nodes in S6 and S7 Figs). We calculated the edges between nodes from three different sources. The first source is the genie3 inference algorithm as presented in (88) and implemented in the *genie3* package in R (documentation at https://bioconductor.org/packages/release/bioc/html/GENIE3.html). This algorithm infers relations between nodes by creating a random forest classifier for each of the genes in the sample and the rest of the genes are used as variables. The algorithm then estimates the relevance of each gene for the classification of the target gene and based on this defines a weight that represent the plausibility of a connection between the two nodes. This value is not a statistical test and cannot be interpreted as such but it constitutes a ranking of the possible interactions between samples. For the CC6 network the threshold value was set to 0.45 and for CC1 the threshold value was set to 0.1. The interactions obtained by this method are marked in the network Figs (S10 and S7 Figs) as dashed lines. Also, we estimate the sign of the interactions (activating or inhibitory) by calculation a partial correlation between samples. This was done using the *pcor* function of the *ppcor* package in R (89). Because of the low sample number, the partial correlations do not reach statistical significance, for that reason we take a conservative approach and set as inhibitory interactions, only those that are lower than −0.3. The other two methods used are data base search based. We searched the string (90) and the biogrid (91) databases. The string database search was made directly on the string database website using the multiprotein search function. The results were pruned to show only experimental, databases or co-expression relations between nodes. We exported the results as a tab delimited table and use it as directed connections for our network. Finally, we consulted the biogrid data base by downloading the *BIOGRID-ALL-4.2.191* interactions file and searching in R for the interactions that involved proteins or genes in our network. Interaction files are available from our website (https://www.biozentrum.uni-wuerzburg.de/bioinfo/computing/CEACAM/). For the final assembly of the network, any duplicate interactions were removed. With the remaining interactions we assembled the networks using the yEd graphics program (freeware, https://www.yworks.com/products/yed).

### Dynamical modeling of the involved response network considering CEACAM6, CEACAM3 and CEACAM1

We performed simulations of the networks in Fig 5 using Jimena (92). Jimena is a software that enables the dynamical simulation of protein networks by modeling the Boolean state of nodes as a continuous hill function (92). This method is recognized to result in validated Boolean and semiquantitative models for systems biology in infections for transcriptome data (93–95). The nodes can have continuous activation values that range from zero to one. The sign of the connections remains as activating or inhibitory. For our simulations we created an extra node to stimulate the network and analyze its response to that stimulus. The extra node created is called PATHOGEN (for the pathogen *C. albicans*) and has an activating connection to each of the CEACAM nodes in the study (CEACAM1, CEACAM3 and CEACAM6). Also, to link the CEACAM receptors to the rest of the network we set interactions from the CEACAM1, CEACAM3 and CEACAM6 nodes to the most two differentially expressed genes in our RNA-seq analysis (IL6 and ACOD1). For CEACAM3, only a connection to *IL6* was established since ACOD1 was not differentially expressed in that case.

### Data availability

Sequencing data are available under the following link: http://www.ncbi.nlm.nih.gov/bioproject/681392. All other data mentioned including the systems biological models and experimental data are available from the manuscript files and its supplementary data.

### Statistics

Statistical analysis of all cell-based assays was performed using One-Way Annova with Bonferroni post-test. Non-linear data (relative fluorescence units) were logarithmized before the analysis.

### Funding Disclosure

This work was supported by the German Research Foundation (Collaborative Research Center/Transregio 124 – FungiNet —Pathogenic fungi and their human host: Networks of Interaction, DFG project number 210879364, Projects B1, B2, and A5) to TD, MD, and HS, and ProChance 2018 – Programmlinie A1 (Förderkennzeichen 2.11.3-A1/218-01) to EK. The funders had no role in study design, data collection and interpretation, or the decision to submit the work for publication.

## Acknowledgement

We thank Simone Tänzer, Moira Stark, Birgit Maranca-Hüwel, and Bärbel Gobs-Hevelke for their excellent technical support.

## Supplemental Figure Legends

**S1 Fig: No alterations of *C. albicans* binding by human neutrophils in response to anti-CEACAM antibodies.** Human neutrophils were stained for CD11b and left untreated or were incubated with 10 µg/ml isotype control antibody (clone MOPC-21), or monoclonal antibodies B3-17 (CC1), 308/3-3 (CC3), or 1H7-4B (CC6) for 45 min and were consecutively incubated with live, APC-labeled *C. albicans* yeast cells (MOI 1) for 20 min. Cells were then analyzed by flow cytometry for the percentage of *C. albicans*-bound neutrophils. The graphs show the percentage of *C. albicans*-bound granulocytes from two independent experiments.

**S2 Fig: No alterations of basic and *C. albicans*-induced ROS production by human neutrophils in response to anti-CEACAM antibodies.** Human neutrophils were stained for CD11b and left untreated or were incubated with 10 µg/ml isotype control antibody (clone MOPC-21), or monoclonal antibodies B3-17 (CC1), 308/3-3 (CC3), or 1H7-4B (CC6) for 45 min and were consecutively incubated with or without live, APC-labeled *C. albicans* yeast cells (MOI 1) for 20 min, respectively. Cells were then analyzed by flow cytometry for the ROS production by DHR123. The graph shows the logarithm of the fluorescent signals of the DHR123 dye and the respective means from two independent experiments.

**S3 Fig: No alterations of *C. albicans* killing by human neutrophils in response to anti-CEACAM antibodies.** 2×10^5^ human neutrophils (2×10^6^/ml) were left untreated or were incubated with 10 µg/ml isotype control antibody (clone MOPC-21), or monoclonal antibodies B3-17 (CC1), 308/3-3 (CC3), or 1H7-4B (CC6) for 45 min and were consecutively incubated with live *C. albicans* yeast cells (MOI 1) for 30 min. Viable *C. albicans* cells were quantified by XTT assay (different concentrations of *C. albicans* alone served as standard, see Materials and Methods section). The graph shows mean and SD from three independent experiments. Statistical analysis was performed by Repeated Measure ANOVA and showed no significant differences.

**S4 Fig: Principal component analysis (PCA) of all sequenced samples.** In two independent experiments, human neutrophils were left untreated or were incubated with 10 µg/ml isotype control antibody (clone MOPC-21), or monoclonal antibodies B3-17 (CC1), 308/3-3 (CC3), or 1H7-4B (CC6) for 45 min and were consecutively incubated with or without live *C. albicans* yeast cells (MOI 1) for 2 h; mRNA was extracted and analyzed by sequencing. Samples analyzed here were also used for analyses presented in Figs 2, 3, 4, 6 and 7, and in S5to S8 Figs, as well as in S1 to S10 Tables. Legend: pre-treatments (45 min): untreated/without antibody (UT, black), IgG istotype control antibody-treated (Iso, grey), B3-17-treated (CC1, red), 308/3-3-treated (CC3, green); 1H7-4B-treated (CC6, blue); stimulations (2 h): without further stimulation/unstimulated (US, square symbols), with *C. albicans* stimulation (+Ca, round symbols).

**S5 Fig: Altered neutrophil transcription by priming with anti-CEACAM antibodies in absence of *C. albicans*.** In two independent experiments, human neutrophils were left untreated or were incubated with 10 µg/ml isotype control antibody (clone MOPC-21), or monoclonal antibodies B3-17 (CC1), 308/3-3 (CC3), or 1H7-4B (CC6) for 45 min and were consecutively incubated with or without live *C. albicans* yeast cells (MOI 1) for 2 h; mRNA was extracted and analyzed by sequencing. Differentially expressed genes (DEGs) from different comparisons (untreated without C.a. vs. untreated with C.a., IgG-treated without C.a. vs. IgG-treated with C.a., IgG-treated without C.a. vs. CC1-treated without C.a., IgG-treated without C.a. vs. CC3-treated without C.a., and IgG-treated without C.a. vs. CC6-treated without C.a.) are displayed in the heat map. The latter three comparisons display the effect of the respective anti-CEACAM antibody treatment alone (in absence of *C. albicans*). Note that only significantly regulated genes with an adjusted p-value <0.05 and a fold-change of at least ±2 in one of the anti-CEACAM-treated samples in absence of *C. albicans* versus IgG-treated treated samples in absence of *C. albicans* were included. Each row was normalized (mean = 0) and scaled (standard deviation = 1). Which CEACAM treatment(s) altered the respective gene is color-coded in the leftmost column of the heatmap and in the Venn diagram that also shows the total number of altered genes (orange: unique for anti-CEACAM1 treatment in absence of *C. albicans*, green: unique for anti-CEACAM6 treatment in absence of *C. albicans*, purple: shared by anti-CEACAM1 and anti-CEACAM6 treatment in absence of *C. albicans*, blue: shared by all three anti-CEACAM treatments in absence of *C. albicans*). Please refer to S4 Fig for a principal component analysis of all samples and to S1 and S2 Tables and S6 to S8 Tables for details on DEGs, as well as complete lists of transcripts of the comparisons displayed in the heat map.

**S6 Fig: Network analysis of transcriptional data from anti-CEACAM6-treated neutrophils in presence of *C. albicans*.** Signaling network assembled from the DEGs in CC6-treated neutrophils in presence of *C. albicans* stimulation versus IgG-treated neutrophils in presence of *C. albicans* stimulation (blue and red nodes, data from Table S5) and known interactors of CEACAM family receptors (gray nodes). Interactions are inferred from the RNA-seq samples (dashed lines) or obtained from interaction databases (solid lines). Nodes belonging to selected significantly enriched GO terms important to neutrophil responses are clustered and framed. Refer to S11 Table for complete list of enriched GO terms. Note that this GO term enrichment analysis is independent from the GO term enrichment analysis based on transcriptional data only presented in Fig 3.

**S7 Fig: Network analysis of transcriptional data from anti-CEACAM1-treated neutrophils in presence of *C. albicans*.** Signaling network assembled from the DEGs in CC1-treated neutrophils in presence of *C. albicans* stimulation (blue and red nodes, S3 Table) and known interactors of CEACAM family receptors (gray nodes). Interactions are inferred from the RNA-seq samples (dashed lines) or obtained from interaction databases (solid lines). Nodes belonging to selected significantly enriched GO terms (cellular functions) are clustered and framed. Note that programmed cell death was not among the significantly enriched GO terms when analyzed as described for S6 Fig.

**S8 Fig. Result curves of dynamical simulations of CEACAM1- and CEACAM6-mediated effects on the *C. albicans*-induced reactions of human neutrophils.** The whole networks around CEACAM1 (S7 Fig) and CEACAM6 (S6 Fig) and each protein activation pathway was modelled and considered in the dynamical simulation of the pathogen stimulation effects. For the simulations the extra node “PATHOGEN” (for *C. albicans*) was added to the network; this node is activating and linked to the CEACAM1 node (upper panel) and the CEACAM6 node (lower panel), respectively. The information flow was modeled using Hill functions (see Materials and Methods section for details). Selected trajectories are presented. Note the absence of caspase induction (CASP1 and CASP8) in the CEACAM1-based simulation (upper panel). Note that *IL1B* was not significant for the CEACAM1 simulation (upper panel) and is therefore not represented here.

## Supplemental Tables

**S1 Table: Gene expression analysis_no antibody-plus Ca versus no antibody-unstimulated.** The table displays the effect of *C. albicans* on the neutrophil transcriptional response after 2 h in absence of antibody treatment and shows the data for all detected transcripts (sheet 1) and for DEGs only (>± 2-fold expression and adjusted p-value < 0.05, sheet 2), respectively.

**S2 Table: Gene expression analysis_IgG-plus Ca versus IgG-unstimulated.** The table displays the effect of *C. albicans* on the neutrophil transcriptional response after 2 h in presence of IgG1 control antibody treatment and shows the data for all detected transcripts (sheet 1) and for DEGs only (>± 2-fold expression and adjusted p-value < 0.05, sheet 2), respectively.

**S3 Table: Gene expression analysis_CC1-plus Ca versus IgG-plus Ca.** The table displays the effect of the anti-CEACAM1-antibody treatment on the neutrophil transcriptional response on top of the effect of the *C. albicans* stimulation for 2 h, and shows the data for all detected transcripts (sheet 1) and for DEGs only (>± 2-fold expression and adjusted p-value < 0.05, sheet 2), respectively.

**S4 Table: Gene expression analysis_CC3-plus Ca versus IgG-plus Ca.** The table displays the effect of the anti-CEACAM3-antibody treatment on the neutrophil transcriptional response on top of the effect of the *C. albicans* stimulation for 2 h, and shows the data for all detected transcripts (sheet 1) and for DEGs only (>± 2-fold expression and adjusted p-value < 0.05, sheet 2), respectively.

**S5 Table: Gene expression analysis_CC6-plus Ca versus IgG-plus Ca.** The table displays the effect of the anti-CEACAM6-antibody treatment on the neutrophil transcriptional response on top of the effect of the *C. albicans* stimulation for 2 h, and shows the data for all detected transcripts (sheet 1) and for DEGs only (>± 2-fold expression and adjusted p-value < 0.05, sheet 2), respectively.

**S6 Table: Gene expression analysis_CC1-unstimulated versus IgG-unstimulated.** The table displays the effect of the anti-CEACAM1-antibody treatment alone on the neutrophil transcriptional response (in the absence of *C. albicans* stimulation), and shows the data for all detected transcripts (sheet 1) and for DEGs only (>± 2-fold expression and adjusted p-value < 0.05, sheet 2), respectively.

**S7 Table: Gene expression analysis_CC3-unstimulated versus IgG-unstimulated.** The table displays the effect of the anti-CEACAM3-antibody treatment alone on the neutrophil transcriptional response (in the absence of *C. albicans* stimulation), and shows the data for all detected transcripts (sheet 1) and for DEGs only (>± 2-fold expression and adjusted p-value < 0.05, sheet 2), respectively.

**S8 Table: Gene expression analysis_CC6-unstimulated versus IgG-unstimulated.** The table displays the effect of the anti-CEACAM6-antibody treatment alone on the neutrophil transcriptional response (in the absence of *C. albicans* stimulation), and shows the data for all detected transcripts (sheet 1) and for DEGs only (>± 2-fold expression and adjusted p-value < 0.05, sheet 2), respectively.

**S9 Table: Enriched GO terms_complete lists.** The table displays the complete lists of significantly enriched GO terms from the following comparisons: IgG/with Ca versus IgG/no Ca (sheet 1), CC1/with Ca versus IgG/with Ca (sheet 2), CC6/with Ca versus IgG/with Ca (sheet 3, see also Fig 3), CC1/no Ca versus IgG/no Ca (sheet 4), CC6/no Ca versus IgG/no Ca (sheet 5). DEGs from the respective comparison were analyzed for enriched GO terms and filtered for test values <0.01 in three different statistical tests (see Materials and Methods section for details).

**S10 Table: Enriched KEGG pathways-network analysis-CC6 plus Ca vs IgG plus Ca_complete list.** The table displays the complete lists of significantly enriched KEGG pathways after anti-CEACAM6 antibody treatment in presence of *C. albicans* stimulation (compared to IgG treatment in presence of *C. albicans* stimulation) from the network analysis displayed in Fig 4 (based on S5 Table, adjusted p-value <0.05).

**S11 Table: Enriched GO terms-network analysis-CC6 plus Ca vs IgG plus Ca_complete list.** The table displays the complete lists of significantly enriched GO terms (cellular functions) after anti-CEACAM6 antibody treatment in presence of *C. albicans* stimulation (compared to IgG treatment in presence of *C. albicans* stimulation) from the network analysis displayed in S6 Fig (based on S5 Table, classic p-value <0.01).

## References

1. Kullberg BJ, Arendrup MC. Invasive Candidiasis. N Engl J Med. 2015;373(15):1445–56.

2. Sonego F, Castanheira FV, Ferreira RG, Kanashiro A, Leite CA, Nascimento DC, et al. Paradoxical Roles of the Neutrophil in Sepsis: Protective and Deleterious. Front Immunol. 2016;7:155.

3. Tamassia N, Bianchetto-Aguilera F, Arruda-Silva F, Gardiman E, Gasperini S, Calzetti F, et al. Cytokine production by human neutrophils: Revisiting the “dark side of the moon”. Eur J Clin Invest. 2018;48 Suppl 2:e12952.

4. Metzemaekers M, Gouwy M, Proost P. Neutrophil chemoattractant receptors in health and disease: double-edged swords. Cell Mol Immunol. 2020;17(5):433–50.

5. Schauer AE, Klassert TE, von Lachner C, Riebold D, Schneeweiss A, Stock M, et al. IL-37 Causes Excessive Inflammation and Tissue Damage in Murine Pneumococcal Pneumonia. J Innate Immun. 2017;9(4):403–18.

6. Azcutia V, Parkos CA, Brazil JC. Role of negative regulation of immune signaling pathways in neutrophil function. J Leukoc Biol. 2017.

7. Sadarangani M, Pollard AJ, Gray-Owen SD. Opa proteins and CEACAMs: pathways of immune engagement for pathogenic Neisseria. FEMS Microbiol Rev. 2011;35(3):498–514.

8. Singer BB, Opp L, Heinrich A, Schreiber F, Binding-Liermann R, Berrocal-Almanza LC, et al. Soluble CEACAM8 interacts with CEACAM1 inhibiting TLR2-triggered immune responses. PLoS One. 2014;9(4):e94106.

9. Slevogt H, Seybold J, Tiwari KN, Hocke AC, Jonatat C, Dietel S, et al. Moraxella catarrhalis is internalized in respiratory epithelial cells by a trigger-like mechanism and initiates a TLR2- and partly NOD1-dependent inflammatory immune response. Cell Microbiol. 2007;9(3):694–707.

10. Lu R, Pan H, Shively JE. CEACAM1 negatively regulates IL-1beta production in LPS activated neutrophils by recruiting SHP-1 to a SYK-TLR4-CEACAM1 complex. PLoS Pathog. 2012;8(4):e1002597.

11. Gray-Owen SD, Blumberg RS. CEACAM1: contact-dependent control of immunity. Nat Rev Immunol. 2006;6(6):433–46.

12. Klaile E, Muller MM, Schafer MR, Clauder AK, Feer S, Heyl KA, et al. Binding of Candida albicans to Human CEACAM1 and CEACAM6 Modulates the Inflammatory Response of Intestinal Epithelial Cells. mBio. 2017;8(2).

13. Singer BB, Scheffrahn I, Heymann R, Sigmundsson K, Kammerer R, Obrink B. Carcinoembryonic antigen-related cell adhesion molecule 1 expression and signaling in human, mouse, and rat leukocytes: evidence for replacement of the short cytoplasmic domain isoform by glycosylphosphatidylinositol-linked proteins in human leukocytes. J Immunol. 2002;168(10):5139–46.

14. Tchoupa AK, Schuhmacher T, Hauck CR. Signaling by epithelial members of the CEACAM family - mucosal docking sites for pathogenic bacteria. Cell Commun Signal. 2014;12:27.

15. Helfrich I, Singer BB. Size Matters: The Functional Role of the CEACAM1 Isoform Signature and Its Impact for NK Cell-Mediated Killing in Melanoma. Cancers (Basel). 2019;11(3).

16. Muller MM, Singer BB, Klaile E, Obrink B, Lucka L. Transmembrane CEACAM1 affects integrin-dependent signaling and regulates extracellular matrix protein-specific morphology and migration of endothelial cells. Blood. 2005;105(10):3925–34.

17. Javaheri A, Kruse T, Moonens K, Mejias-Luque R, Debraekeleer A, Asche CI, et al. Helicobacter pylori adhesin HopQ engages in a virulence-enhancing interaction with human CEACAMs. Nat Microbiol. 2016;2:16189.

18. Slevogt H, Zabel S, Opitz B, Hocke A, Eitel J, N’Guessan P D, et al. CEACAM1 inhibits Toll-like receptor 2-triggered antibacterial responses of human pulmonary epithelial cells. Nat Immunol. 2008;9(11):1270–8.

19. Kammerer R, Hahn S, Singer BB, Luo JS, von Kleist S. Biliary glycoprotein (CD66a), a cell adhesion molecule of the immunoglobulin superfamily, on human lymphocytes: structure, expression and involvement in T cell activation. Eur J Immunol. 1998;28(11):3664–74.

20. Singer BB, Klaile E, Scheffrahn I, Muller MM, Kammerer R, Reutter W, et al. CEACAM1 (CD66a) mediates delay of spontaneous and Fas ligand-induced apoptosis in granulocytes. Eur J Immunol. 2005;35(6):1949–59.

21. Khairnar V, Duhan V, Maney SK, Honke N, Shaabani N, Pandyra AA, et al. CEACAM1 induces B-cell survival and is essential for protective antiviral antibody production. Nat Commun. 2015;6:6217.

22. Khairnar V, Duhan V, Patil AM, Zhou F, Bhat H, Thoens C, et al. CEACAM1 promotes CD8(+) T cell responses and improves control of a chronic viral infection. Nat Commun. 2018;9(1):2561.

23. Bonsignore P, Kuiper JWP, Adrian J, Goob G, Hauck CR. CEACAM3-A Prim(at)e Invention for Opsonin-Independent Phagocytosis of Bacteria. Front Immunol. 2019;10:3160.

24. Sintsova A, Guo CX, Sarantis H, Mak TW, Glogauer M, Gray-Owen SD. Bcl10 synergistically links CEACAM3 and TLR-dependent inflammatory signalling. Cell Microbiol. 2018;20(1).

25. Heinrich A, Heyl KA, Klaile E, Muller MM, Klassert TE, Wiessner A, et al. Moraxella catarrhalis induces CEACAM3-Syk-CARD9-dependent activation of human granulocytes. Cell Microbiol. 2016;18(11):1570–82.

26. Gutbier B, Fischer K, Doehn JM, von Lachner C, Herr C, Klaile E, et al. Moraxella catarrhalis induces an immune response in the murine lung that is independent of human CEACAM5 expression and long-term smoke exposure. Am J Physiol Lung Cell Mol Physiol. 2015;309(3):L250–61.

27. Pils S, Kopp K, Peterson L, Delgado Tascon J, Nyffenegger-Jann NJ, Hauck CR. The adaptor molecule Nck localizes the WAVE complex to promote actin polymerization during CEACAM3-mediated phagocytosis of bacteria. PLoS One. 2012;7(3):e32808.

28. Sarantis H, Gray-Owen SD. Defining the roles of human carcinoembryonic antigen-related cellular adhesion molecules during neutrophil responses to Neisseria gonorrhoeae. Infect Immun. 2012;80(1):345–58.

29. Tegtmeyer N, Ghete TD, Schmitt V, Remmerbach T, Cortes MCC, Bondoc EM, et al. Type IV secretion of Helicobacter pylori CagA into oral epithelial cells is prevented by the absence of CEACAM receptor expression. Gut Pathog. 2020;12:25.

30. Tegtmeyer N, Harrer A, Schmitt V, Singer BB, Backert S. Expression of CEACAM1 or CEACAM5 in AZ-521 cells restores the type IV secretion deficiency for translocation of CagA by Helicobacter pylori. Cell Microbiol. 2019;21(1):e12965.

31. Behrens IK, Busch B, Ishikawa-Ankerhold H, Palamides P, Shively JE, Stanners C, et al. The HopQ-CEACAM Interaction Controls CagA Translocation, Phosphorylation, and Phagocytosis of Helicobacter pylori in Neutrophils. mBio. 2020;11(1).

32. Pan H, Shively JE. Carcinoembryonic antigen-related cell adhesion molecule-1 regulates granulopoiesis by inhibition of granulocyte colony-stimulating factor receptor. Immunity. 2010;33(4):620–31.

33. Skubitz KM, Campbell KD, Skubitz AP. CD66a, CD66b, CD66c, and CD66d each independently stimulate neutrophils. J Leukoc Biol. 1996;60(1):106-17.

34. Skubitz KM, Skubitz AP. Interdependency of CEACAM-1, −3, −6, and −8 induced human neutrophil adhesion to endothelial cells. J Transl Med. 2008;6:78.

35. Saalbach A, Arnhold J, Lessig J, Simon JC, Anderegg U. Human Thy-1 induces secretion of matrix metalloproteinase-9 and CXCL8 from human neutrophils. Eur J Immunol. 2008;38(5):1391–403.

36. Pellme S, Morgelin M, Tapper H, Mellqvist UH, Dahlgren C, Karlsson A. Localization of human neutrophil interleukin-8 (CXCL-8) to organelle(s) distinct from the classical granules and secretory vesicles. J Leukoc Biol. 2006;79(3):564–73.

37. Hatanaka E, Monteagudo PT, Marrocos MS, Campa A. Neutrophils and monocytes as potentially important sources of proinflammatory cytokines in diabetes. Clin Exp Immunol. 2006;146(3):443–7.

38. Hidalgo MA, Carretta MD, Teuber SE, Zarate C, Carcamo L, Concha, II, et al. fMLP-Induced IL-8 Release Is Dependent on NADPH Oxidase in Human Neutrophils. J Immunol Res. 2015;2015:120348.

39. Klaile E, Muller MM, Zubiria-Barrera C, Brehme S, Klassert TE, Stock M, et al. Unaltered Fungal Burden and Lethality in Human CEACAM1-Transgenic Mice During Candida albicans Dissemination and Systemic Infection. Front Microbiol. 2019;10:2703.

40. Niemiec MJ, Grumaz C, Ermert D, Desel C, Shankar M, Lopes JP, et al. Dual transcriptome of the immediate neutrophil and Candida albicans interplay. BMC Genomics. 2017;18(1):696.

41. Philip NH, Dillon CP, Snyder AG, Fitzgerald P, Wynosky-Dolfi MA, Zwack EE, et al. Caspase-8 mediates caspase-1 processing and innate immune defense in response to bacterial blockade of NF-kappaB and MAPK signaling. Proc Natl Acad Sci U S A. 2014;111(20):7385–90.

42. Fritsch M, Gunther SD, Schwarzer R, Albert MC, Schorn F, Werthenbach JP, et al. Caspase-8 is the molecular switch for apoptosis, necroptosis and pyroptosis. Nature. 2019;575(7784):683-7.

43. Schirbel A, Rebert N, Sadler T, West G, Rieder F, Wagener C, et al. Mutual Regulation of TLR/NLR and CEACAM1 in the Intestinal Microvasculature: Implications for IBD Pathogenesis and Therapy. Inflamm Bowel Dis. 2019;25(2):294–305.

44. Bourgeois C, Kuchler K. Fungal pathogens-a sweet and sour treat for toll-like receptors. Front Cell Infect Microbiol. 2012;2:142.

45. Sarantis H, Gray-Owen SD. The specific innate immune receptor CEACAM3 triggers neutrophil bactericidal activities via a Syk kinase-dependent pathway. Cell Microbiol. 2007;9(9):2167–80.

46. Sintsova A, Sarantis H, Islam EA, Sun CX, Amin M, Chan CH, et al. Global analysis of neutrophil responses to Neisseria gonorrhoeae reveals a self-propagating inflammatory program. PLoS Pathog. 2014;10(9):e1004341.

47. Schmitter T, Agerer F, Peterson L, Munzner P, Hauck CR. Granulocyte CEACAM3 is a phagocytic receptor of the innate immune system that mediates recognition and elimination of human-specific pathogens. J Exp Med. 2004;199(1):35–46.

48. Schmitter T, Pils S, Sakk V, Frank R, Fischer KD, Hauck CR. The granulocyte receptor carcinoembryonic antigen-related cell adhesion molecule 3 (CEACAM3) directly associates with Vav to promote phagocytosis of human pathogens. J Immunol. 2007;178(6):3797–805.

49. Schmitter T, Pils S, Weibel S, Agerer F, Peterson L, Buntru A, et al. Opa proteins of pathogenic neisseriae initiate Src kinase-dependent or lipid raft-mediated uptake via distinct human carcinoembryonic antigen-related cell adhesion molecule isoforms. Infect Immun. 2007;75(8):4116–26.

50. Hauck CR, Lorenzen D, Saas J, Meyer TF. An in vitro-differentiated human cell line as a model system to study the interaction of Neisseria gonorrhoeae with phagocytic cells. Infect Immun. 1997;65(5):1863–9.

51. Zhang Z, La Placa D, Nguyen T, Kujawski M, Le K, Li L, et al. CEACAM1 regulates the IL-6 mediated fever response to LPS through the RP105 receptor in murine monocytes. BMC Immunol. 2019;20(1):7.

52. Slaats J, Ten Oever J, van de Veerdonk FL, Netea MG. IL-1beta/IL-6/CRP and IL-18/ferritin: Distinct Inflammatory Programs in Infections. PLoS Pathog. 2016;12(12):e1005973.

53. Muller MM, Klaile E, Vorontsova O, Singer BB, Obrink B. Homophilic adhesion and CEACAM1-S regulate dimerization of CEACAM1-L and recruitment of SHP-2 and c-Src. J Cell Biol. 2009;187(4):569–81.

54. Singer BB, Scheffrahn I, Obrink B. The tumor growth-inhibiting cell adhesion molecule CEACAM1 (C-CAM) is differently expressed in proliferating and quiescent epithelial cells and regulates cell proliferation. Cancer Res. 2000;60(5):1236–44.

55. Shikotra A, Choy DF, Siddiqui S, Arthur G, Nagarkar DR, Jia G, et al. A CEACAM6-High Airway Neutrophil Phenotype and CEACAM6-High Epithelial Cells Are Features of Severe Asthma. J Immunol. 2017;198(8):3307–17.

56. Aleandri M, Conte MP, Simonetti G, Panella S, Celestino I, Checconi P, et al. Influenza A virus infection of intestinal epithelial cells enhances the adhesion ability of Crohn’s disease associated Escherichia coli strains. PLoS One. 2015;10(2):e0117005.

57. Klaile E, Klassert TE, Scheffrahn I, Muller MM, Heinrich A, Heyl KA, et al. Carcinoembryonic antigen (CEA)-related cell adhesion molecules are co-expressed in the human lung and their expression can be modulated in bronchial epithelial cells by non-typable Haemophilus influenzae, Moraxella catarrhalis, TLR3, and type I and II interferons. Respir Res. 2013;14:85.

58. Negroni A, Costanzo M, Vitali R, Superti F, Bertuccini L, Tinari A, et al. Characterization of adherent-invasive Escherichia coli isolated from pediatric patients with inflammatory bowel disease. Inflamm Bowel Dis. 2012;18(5):913–24.

59. Barnich N, Carvalho FA, Glasser AL, Darcha C, Jantscheff P, Allez M, et al. CEACAM6 acts as a receptor for adherent-invasive E. coli, supporting ileal mucosa colonization in Crohn disease. J Clin Invest. 2007;117(6):1566–74.

60. Carvalho FA, Barnich N, Sivignon A, Darcha C, Chan CH, Stanners CP, et al. Crohn’s disease adherent-invasive Escherichia coli colonize and induce strong gut inflammation in transgenic mice expressing human CEACAM. J Exp Med. 2009;206(10):2179–89.

61. Han ZM, Huang HM, Sun YW. Effect of CEACAM-1 knockdown in human colorectal cancer cells. Oncol Lett. 2018;16(2):1622–6.

62. N’Guessan PD, Vigelahn M, Bachmann S, Zabel S, Opitz B, Schmeck B, et al. The UspA1 protein of Moraxella catarrhalis induces CEACAM-1-dependent apoptosis in alveolar epithelial cells. J Infect Dis. 2007;195(11):1651–60.

63. Cameron S, de Long LM, Hazar-Rethinam M, Topkas E, Endo-Munoz L, Cumming A, et al. Focal overexpression of CEACAM6 contributes to enhanced tumourigenesis in head and neck cancer via suppression of apoptosis. Mol Cancer. 2012;11:74.

64. Chan CH, Camacho-Leal P, Stanners CP. Colorectal hyperplasia and dysplasia due to human carcinoembryonic antigen (CEA) family member expression in transgenic mice. PLoS One. 2007;2(12):e1353.

65. Duxbury MS, Matros E, Ito H, Zinner MJ, Ashley SW, Whang EE. Systemic siRNA-mediated gene silencing: a new approach to targeted therapy of cancer. Ann Surg. 2004;240(4):667–74; discussion 75-6.

66. Gaur P, Ranjan P, Sharma S, Patel JR, Bowzard JB, Rahman SK, et al. Influenza A virus neuraminidase protein enhances cell survival through interaction with carcinoembryonic antigen-related cell adhesion molecule 6 (CEACAM6) protein. J Biol Chem. 2012;287(18):15109–17.

67. Kolla V, Gonzales LW, Bailey NA, Wang P, Angampalli S, Godinez MH, et al. Carcinoembryonic cell adhesion molecule 6 in human lung: regulated expression of a multifunctional type II cell protein. Am J Physiol Lung Cell Mol Physiol. 2009;296(6):L1019–30.

68. Riley CJ, Engelhardt KP, Saldanha JW, Qi W, Cooke LS, Zhu Y, et al. Design and activity of a murine and humanized anti-CEACAM6 single-chain variable fragment in the treatment of pancreatic cancer. Cancer Res. 2009;69(5):1933–40.

69. Tian C, Zhang B, Ge C. Effect of CEACAM6 silencing on the biological behavior of human gallbladder cancer cells. Oncol Lett. 2020;20(3):2677–88.

70. Kanderova V, Hrusak O, Kalina T. Aberrantly expressed CEACAM6 is involved in the signaling leading to apoptosis of acute lymphoblastic leukemia cells. Exp Hematol. 2010;38(8):653–60 e1.

71. Klaile E, Vorontsova O, Sigmundsson K, Muller MM, Singer BB, Ofverstedt LG, et al. The CEACAM1 N-terminal Ig domain mediates cis- and trans-binding and is essential for allosteric rearrangements of CEACAM1 microclusters. J Cell Biol. 2009;187(4):553–67.

72. Klassert TE, Brauer J, Holzer M, Stock M, Riege K, Zubiria-Barrera C, et al. Differential Effects of Vitamins A and D on the Transcriptional Landscape of Human Monocytes during Infection. Sci Rep. 2017;7:40599.

73. Klassert TE, Hanisch A, Brauer J, Klaile E, Heyl KA, Mansour MK, et al. Modulatory role of vitamin A on the Candida albicans-induced immune response in human monocytes. Med Microbiol Immunol. 2014;203(6):415–24.

74. Afgan E, Baker D, Batut B, van den Beek M, Bouvier D, Cech M, et al. The Galaxy platform for accessible, reproducible and collaborative biomedical analyses: 2018 update. Nucleic Acids Res. 2018;46(W1):W537–W44.

75. Love MI, Huber W, Anders S. Moderated estimation of fold change and dispersion for RNA-seq data with DESeq2. Genome Biol. 2014;15(12):550.

76. Robinson MD, McCarthy DJ, Smyth GK. edgeR: a Bioconductor package for differential expression analysis of digital gene expression data. Bioinformatics. 2010;26(1):139–40.

77. Alexa A, Rahnenfuhrer J. Enrichment Analysis for Gene Ontology. R package version 2.42.0. https://bioconductor.org/packages/release/bioc/html/topGO.html2020.

78. Supek F, Bosnjak M, Skunca N, Smuc T. REVIGO summarizes and visualizes long lists of gene ontology terms. PLoS One. 2011;6(7): e21800.

79. Remmele CW, Luther CH, Balkenhol J, Dandekar T, Muller T, Dittrich MT. Integrated inference and evaluation of host-fungi interaction networks. Front Microbiol. 2015;6: 764.

80. Orchard S, Kerrien S, Abbani S, Aranda B, Bhate J, Bidwell S, et al. Protein interaction data curation: the International Molecular Exchange (IMEx) consortium. Nat Methods. 2012; 9(4): 345-50.

81. Aranda B, Blankenburg H, Kerrien S, Brinkman FS, Ceol A, Chautard E, et al. PSICQUIC and PSISCORE: accessing and scoring molecular interactions. Nat Methods. 2011; 8(7): 528-9.

82. Beisser D, Klau GW, Dandekar T, Muller T, Dittrich MT. BioNet: an R-Package for the functional analysis of biological networks. Bioinformatics. 2010; 26(8): 1129-30.

83. Dittrich MT, Klau GW, Rosenwald A, Dandekar T, Mulleri T. Identifying functional modules in protein-protein interaction networks: an integrated exact approach. Bioinformatics. 2008; 24(13): i223-31.

84. Shannon P, Markiel A, Ozier O, Baliga NS, Wang JT, Ramage D, et al. Cytoscape: a software environment for integrated models of biomolecular interaction networks. Genome Res. 2003; 13(11): 2498-504.

85. Raudvere U, Kolberg L, Kuzmin I, Arak T, Adler P, Peterson H, et al. g:Profiler: a web server for functional enrichment analysis and conversions of gene lists (2019 update). Nucleic Acids Res. 2019;47(W1):W191–W8.

86. Kanehisa M, Furumichi M, Tanabe M, Sato Y, Morishima K. KEGG: new perspectives on genomes, pathways, diseases and drugs. Nucleic Acids Res. 2017;45(D1):D353–D61.

87. UniProt C. UniProt: a worldwide hub of protein knowledge. Nucleic Acids Res. 2019;47(D1):D506–D15.

88. Huynh-Thu VA, Irrthum A, Wehenkel L, Geurts P. Inferring regulatory networks from expression data using tree-based methods. PLoS One. 2010;5(9).

89. Kim S. ppcor: An R Package for a Fast Calculation to Semi-partial Correlation Coefficients. Commun Stat Appl Methods. 2015;22(6):665–74.

90. Szklarczyk D, Gable AL, Lyon D, Junge A, Wyder S, Huerta-Cepas J, et al. STRING v11: protein-protein association networks with increased coverage, supporting functional discovery in genome-wide experimental datasets. Nucleic Acids Res. 2019;47(D1):D607–D13.

91. Stark C, Breitkreutz BJ, Reguly T, Boucher L, Breitkreutz A, Tyers M. BioGRID: a general repository for interaction datasets. Nucleic Acids Res. 2006;34(Database issue):D535-9.

92. Karl S, Dandekar T. Jimena: efficient computing and system state identification for genetic regulatory networks. BMC Bioinformatics. 2013;14:306.

93. Yang M, Rajeeve K, Rudel T, Dandekar T. Comprehensive Flux Modeling of Chlamydia trachomatis Proteome and qRT-PCR Data Indicate Biphasic Metabolic Differences Between Elementary Bodies and Reticulate Bodies During Infection. Front Microbiol. 2019;10:2350.

94. Srivastava M, Bencurova E, Gupta SK, Weiss E, Loffler J, Dandekar T. Aspergillus fumigatus Challenged by Human Dendritic Cells: Metabolic and Regulatory Pathway Responses Testify a Tight Battle. Front Cell Infect Microbiol. 2019;9:168.

95. Cecil A, Gentschev I, Adelfinger M, Dandekar T, Szalay AA. Vaccinia virus injected human tumors: oncolytic virus efficiency predicted by antigen profiling analysis fitted boolean models. Bioengineered. 2019;10(1):190–6.

