## Supplemental Figures for "Antibody ligation of carcinoembryonic antigen-related cell adhesion molecule 1 (CEACAM1), CEACAM3, and CEACAM6, differentially enhance the cytokine release of human neutrophils in responses to *Candida albicans*"

### S1 Fig

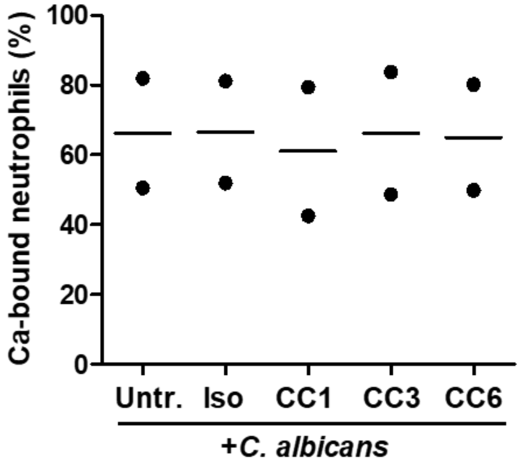


**S1 Fig : No alterations of *C. albicans* bindingby human neutrophils in response to anti-CEACAM antibodies.** Human neutrophils were stained for CD11b and left untreated or were incubated with 10 µg/ml isotype control antibody (clone MOPC-21), or monoclonal antibodies B3-17 (CC1), 308/3-3 (CC3), or 1H7-4B (CC6) for 45 min and were consecutively incubated with live, APC-labeled *C. albicans* yeast cells (MOI 1) for 20 min. Cells were then analyzed by flow cytometry for the percentage of *C. albicans*-bound neutrophils. The graphs show the percentage of *C. albicans*-bound granulocytes from two independent experiments.

### S2 Fig

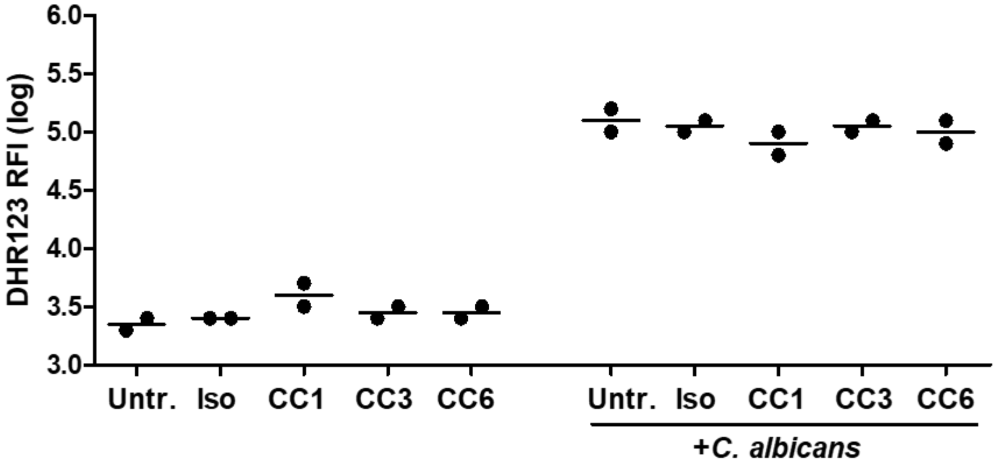


**S2 Fig : No alterations of basic and *C. albicans*-induced ROS production by human neutrophils in response to anti-CEACAM antibodies.** Human neutrophils were stained for CD11b and left untreated or were incubated with 10 µg/ml isotype control antibody (clone MOPC-21), or monoclonal antibodies B3-17 (CC1), 308/3-3 (CC3), or 1H7-4B (CC6) for 45 min and were consecutively incubated with or without live, APC-labeled *C. albicans* yeast cells (MOI 1) for 20 min, respectively. Cells were then analyzed by flow cytometry for the ROS production by DHR123. The graph shows the logarithmized fluorescent signal of the DHR123 dye and the respective means from two independent experiments.

### S3 Fig

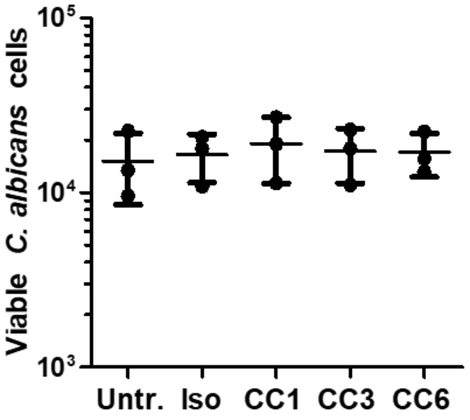


**S3 Fig : No alterations of *C. albicans* killing by human neutrophils in response to anti-CEACAM antibodies.** $2\times10^5$ human neutrophils ($2\times10^6$/ml) were left untreated or were incubated with 10 µg/ml isotype control antibody (clone MOPC-21), or monoclonal antibodies B3-17 (CC1), 308/3-3 (CC3), or 1H7-4B (CC6) for 45 min and were consecutively incubated with live *C. albicans* yeast cells (MOI 1) for 30 min. Viable *C. albicans* cells were quantified by XTT assay (different concentrations of *C. albicans* alone served as standard, see Materials and Methods section). The graph shows mean and SD from three independent experiments. Statistical analysis was performed by Repeated Measure ANOVA and showed no significant differences.

### S4 Fig

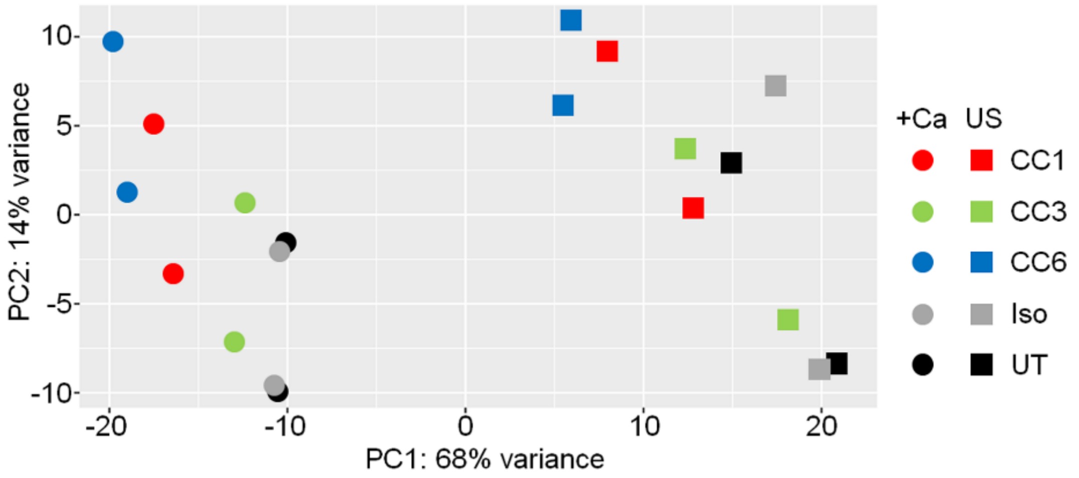


**S4 Fig : Principal component analysis (PCA) of all sequenced samples.** In two independent experiments, human neutrophils were left untreated or were incubated with 10 µg/ml isotype control antibody (clone MOPC-21), or monoclonal antibodies B3-17 (CC1), 308/3-3 (CC3), or 1H7-4B (CC6) for 45 min and were consecutively incubated with or without live *C. albicans* yeast cells (MOI 1) for 2 h; mRNA was extracted and analyzed by sequencing. Samples analyzed here were also used for analyses presented in Figs 2, 3, 4, 6 and 7, and in S5to S12 Figs, as well as in S1 to S10 Tables. Legend: pre-treatments (45 min): untreated/without antibody (UT, black), IgG isotype control antibody-treated (Iso, grey), B3-17-treated (CC1, red), 308/3-3-treated (CC3, green); 1H7-4B-treated (CC6, blue); stimulations (2 h): without further stimulation/unstimulated (US, square symbols), with *C. albicans* stimulation (+Ca, round symbols).

#### S5 Fig

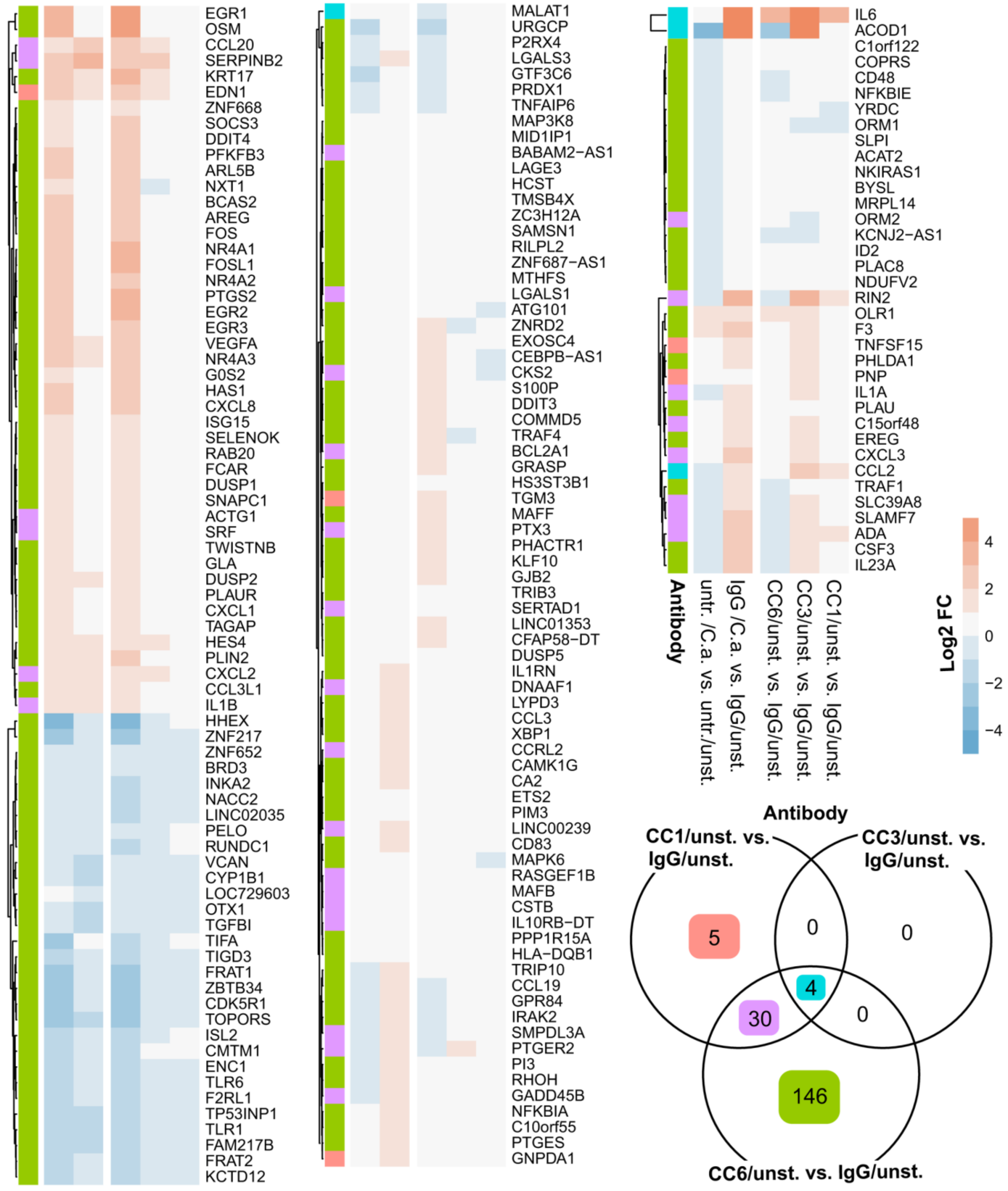

EGR1
OSM
CCL20
SERPINB2
KRT17
EDN1
ZNF668
SOCS3
DDIT4
PFKFB3
ARL5B
NXT1
BCAS2
AREG
FOS
NR4A1
FOSL1
NR4A2
PTGS2
EGR2
EGR3
VEGFA
NR4A3
G0S2
HAS1
CXCL8
ISG15
SELENOK
RAB20
FCAR
DUSP1
SNAPC1
ACTG1
SRF
TWISTNB
GLA
DUSP2
PLAUR
CXCL1
TAGAP
HES4
PLIN2
CXCL2
CCL3L1
IL1B
HHEX
ZNF217
ZNF652
BRD3
INKA2
NACC2
LINC02035
PELO
RUNDC1
VCAN
CYP1B1
LOC729603
OTX1
TGFBI
TIFA
TIGD3
FRAT1
ZBTB34
CDK5R1
TOPORS
ISL2
CMTM1
ENC1
TLR6
F2RL1
TP53INP1
TLR1
FAM217B
FRAT2
KCTD12
MALAT1
URGCP
P2RX4
LGALS3
GTF3C6
PRDX1
TNFAIP6
MAP3K8
MID1IP1
BABAM2−AS1
LAGE3
HCST
TMSB4X
ZC3H12A
SAMSN1
RILPL2
ZNF687−AS1
MTHFS
LGALS1
ATG101
ZNRD2
EXOSC4
CEBPB−AS1
CKS2
S100P
DDIT3
COMMD5
TRAF4
BCL2A1
GRASP
HS3ST3B1
TGM3
MAFF
PTX3
PHACTR1
KLF10
GJB2
TRIB3
SERTAD1
LINC01353
CFAP58−DT
DUSP5
IL1RN
DNAAF1
LYPD3
CCL3
XBP1
CCRL2
CAMK1G
CA2
ETS2
PIM3
LINC00239
CD83
MAPK6
RASGEF1B
MAFB
CSTB
IL10RB−DT
PPP1R15A
HLA−DQB1
TRIP10
CCL19
GPR84
IRAK2
SMPDL3A
PTGER2
PI3
RHOH
GADD45B
NFKBIA
C10orf55
PTGES
GNPDA1
IL6
ACOD1
C1orf122
COPRS
CD48
NFKBIE
YRDC
ORM1
SLPI
ACAT2
NKIRAS1
BYSL
MRPL14
ORM2
KCNJ2−AS1
ID2
PLAC8
NDUFV2
RIN2
OLR1
F3
TNFSF15
PHLDA1
PNP
IL1A
PLAU
C15orf48
EREG
CXCL3
CCL2
TRAF1
SLC39A8
SLAMF7
ADA
CSF3
IL23A
Antibody
untr. /C.a. vs. untr./unst.
IgG /C.a. vs. IgG/unst.
CC6/unst. vs. IgG/unst.
CC3/unst. vs. IgG/unst.
CC1/unst. vs. IgG/unst.
Log2 FC
4
2
0
−2
−4
Antibody
CC1/unst. vs. IgG/unst.
CC3/unst. vs. IgG/unst.
5
0
0
4
30
0
146
CC6/unst. vs. IgG/unst.

**S5 Fig : Altered neutrophil transcription by priming with anti-CEACAM antibodies in absence of *C. albicans*.** In two independent experiments, human neutrophils were left untreated or were incubated with 10 µg/ml isotype control antibody (clone MOPC-21), or monoclonal antibodies B3-17 (CC1), 308/3-3 (CC3), or 1H7-4B (CC6) for 45 min and were consecutively incubated with or without live *C. albicans* yeast cells (MOI 1) for 2 h; mRNA was extracted and analyzed by sequencing. Differentially expressed genes (DEGs) from different comparisons (untreated without C.a. vs. untreated with C.a., IgG-treated without C.a. vs. IgG-treated with C.a., IgG-treated without C.a. vs. CC1-treated without C.a., IgG-treated without C.a. vs. CC3-treated without C.a., and IgG-treated without C.a. vs. CC6-treated without C.a.) are displayed in the heat map. The latter three comparisons display the effect of the respective anti-CEACAM antibody treatment alone (in absence of *C. albicans*). Note that only significantly regulated genes with an adjusted p-value <0.05 and a fold-change of at least ±2 in one of the anti-CEACAM-treated samples in absence of *C. albicans* versus IgG-treated treated samples in absence of *C. albicans* were included. Each row was normalized (mean = 0) and scaled (standard deviation = 1). Which CEACAM treatment(s) altered the respective gene is color-coded in the leftmost column of the heatmap and in the Venn diagram that also shows the total number of altered genes (orange: unique for anti-CEACAM1 treatment in absence of *C. albicans*, green: unique for anti-CEACAM6 treatment in absence of *C. albicans*, purple: shared by anti-CEACAM1 and anti-CEACAM6 treatment in absence of *C. albicans*, blue: shared by all three anti-CEACAM treatments in absence of *C. albicans*). Please refer to S4 Fig for a principal component analysis of all samples and to S1 and S2 Tables and S6 to S8 Tables for details on DEGs, as well as complete lists of transcripts of the comparisons displayed in the heat map.

### S6 Fig

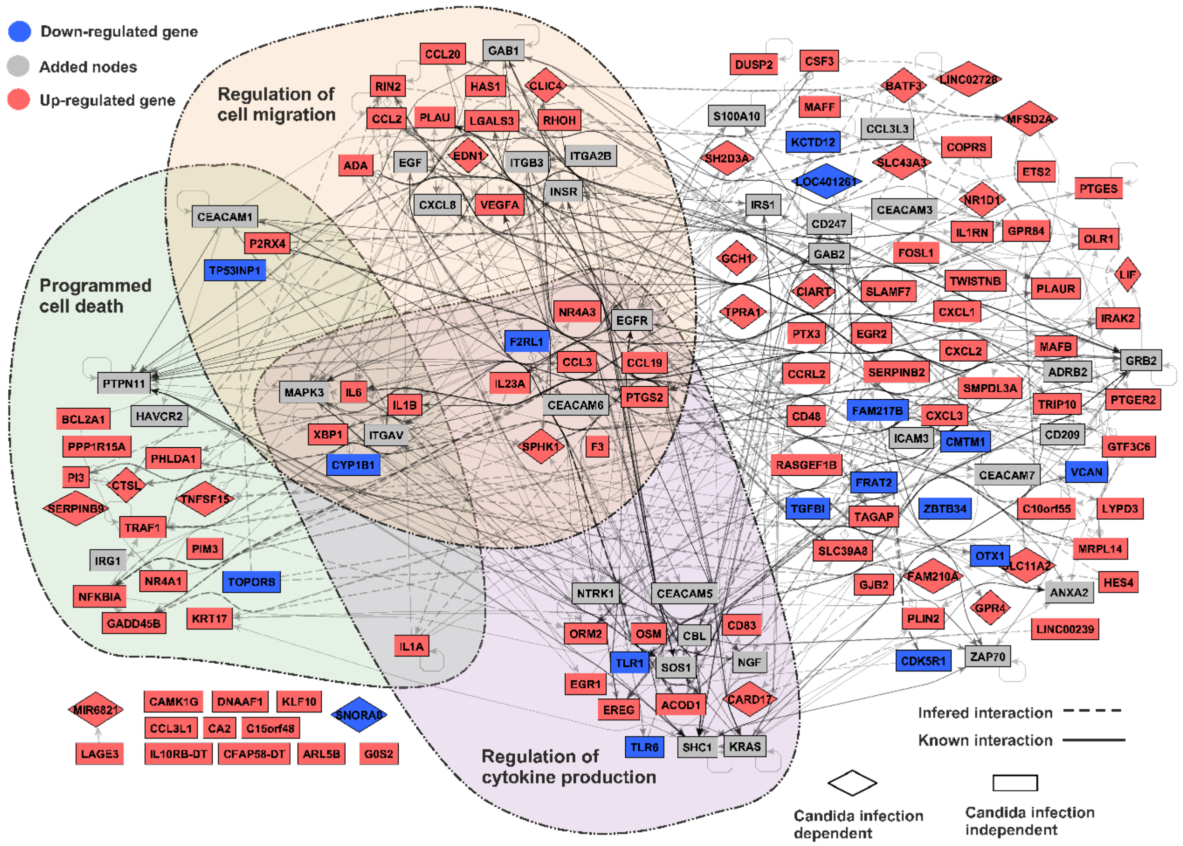


**S6 Fig: Network analysis of transcriptional data from anti-CEACAM6-treated neutrophils in presence of *C. albicans*.** Signaling network assembled from the DEGs in CC6-treated neutrophils in presence of *C. albicans* stimulation versus IgG-treated neutrophils in presence of *C. albicans* stimulation (blue and red nodes, data from Table S5) and known interactors of CEACAM family receptors (gray nodes). Interactions are inferred from the RNA-seq samples (dashed lines) or obtained from interaction databases (solid lines). Nodes belonging to selected significantly enriched GO terms important to neutrophil responses (regulation of cell migration, regulation of cytokine production, programmed cell death) are clustered and framed.

### S7 Fig

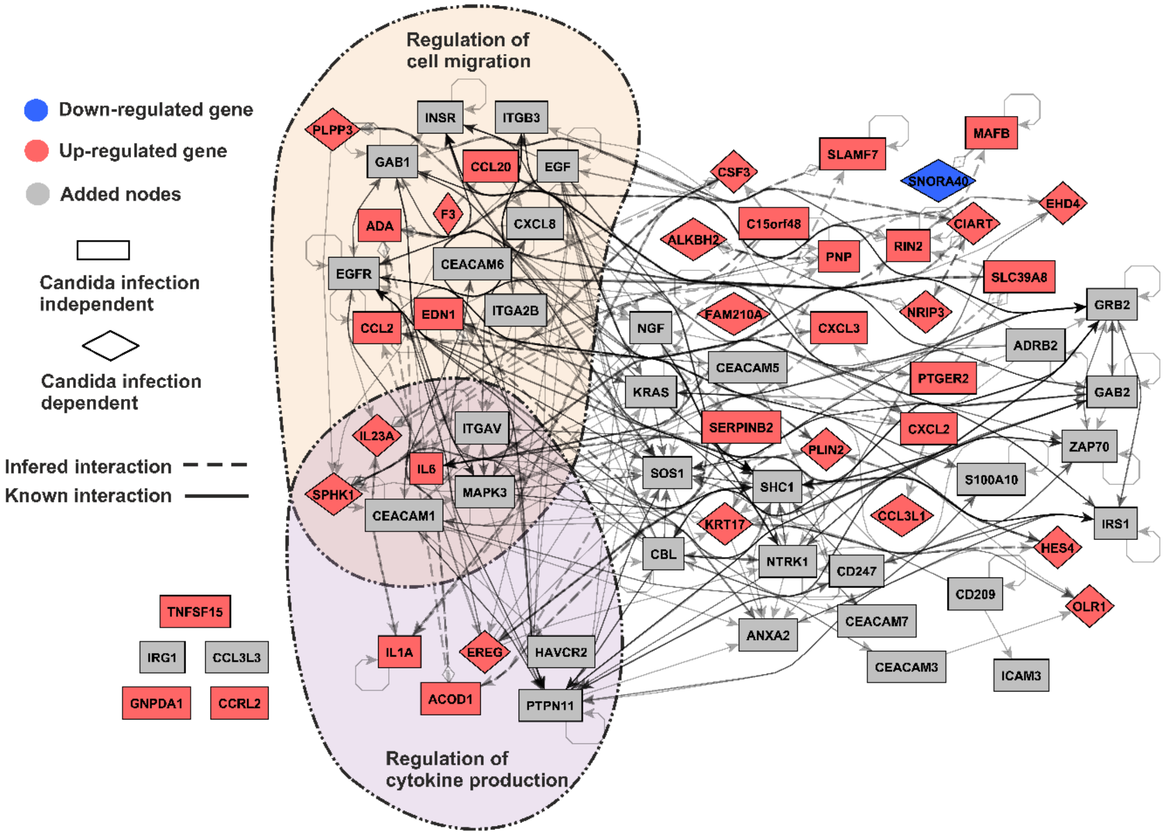


**S7 Fig: Network analysis of transcriptional data from anti-CEACAM1-treated neutrophils in presence of *C. albicans*.** Signaling network assembled from the DEGs in CC1-treated neutrophils in presence of *C. albicans* stimulation versus IgG-treated neutrophils in presence of *C. albicans* stimulation (blue and red nodes, data from S3 Table) and known interactors of CEACAM family receptors (gray nodes). Interactions are inferred from the RNA-seq samples (dashed lines) or obtained from interaction databases (solid lines). Nodes belonging to selected significantly enriched GO terms (cellular functions) are clustered and framed. Note that programmed cell death was not among the significantly enriched GO terms when analyzed as described for S10 Fig.

### S8 Fig

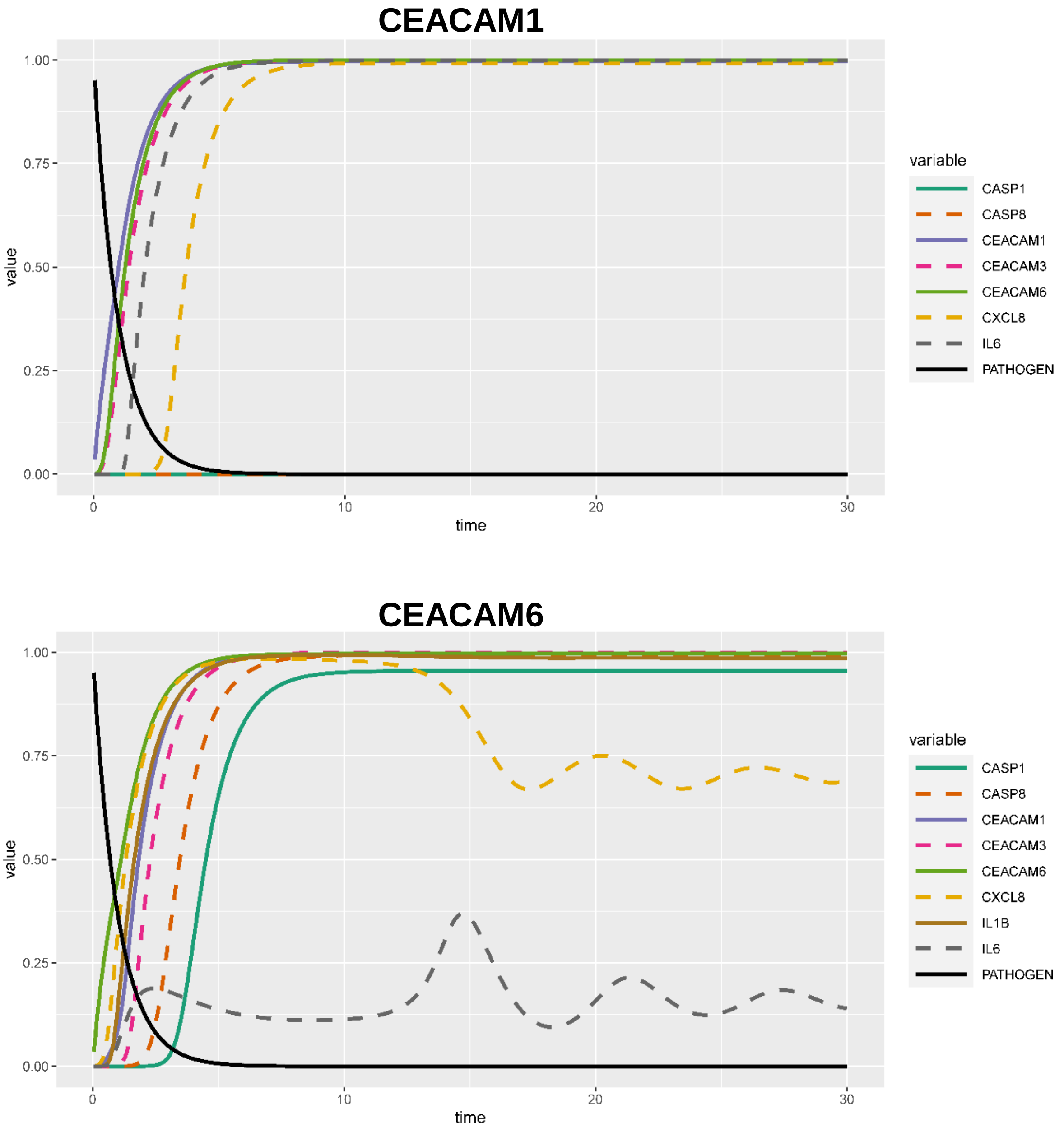


**S8 Fig. Result curves of dynamical simulations of CEACAM1- and CEACAM6-mediated effects on the *C. albicans*-induced reactions of human neutrophils.** The whole networks around CEACAM1 (S11 Fig) and CEACAM6 (S10 Fig) and each protein activation pathway was modelled and considered in the dynamical simulation of the pathogen stimulation effects. For the simulations the extra node "PATHOGEN" (for *C. albicans*) was added to the network; this node is activating and linked to the CEACAM1 node (upper panel) and the CEACAM6 node (lower panel), respectively. The information flow was modeled using Hill functions (see Materials and Methods section for details). Selected trajectories are presented. Note the absence of caspase induction (CASP1 and CASP8) in the CEACAM1-based simulation (upper panel). Note that *IL1B* was not significant for the CEACAM1 simulation (upper panel) and is therefore not represented here.
